# GRACE: Discrete Biological Sequence Optimization with Gradient-Resolved Adaptive Constrained Exploration

**DOI:** 10.64898/2026.11.25.769355

**Authors:** Md Toki Tahmid, Minh Hoang, Mona Singh

**Affiliations:** Princeton University

## Abstract

Machine learning increasingly enables biological sequence redesign for therapeutic and biotechnological applications by optimizing sequences for desired properties. However, improving one property can compromise another: for example, increasing enzyme activity may reduce stability, while enhancing regulatory DNA activity may diminish cell-type specificity. We formulate sequence redesign as a constrained optimization problem guided by learned predictive models, where a reference sequence is mutated to improve a predicted target property while keeping predicted auxiliary properties within specified tolerances of their reference values. To solve this problem, we introduce GRACE (Gradient-Resolved Adaptive Constrained Exploration), a method that optimizes discrete biological sequences using gradients from property predictors via a straight-through estimator. GRACE combines two complementary mechanisms: null-space projection, which removes target-gradient components that interfere with active constraints, and an augmented Lagrangian mechanism, which encourages feasibility after constraint violations. We evaluate GRACE across fourteen diverse tasks, including six protein optimization tasks (spanning activity, stability, and solubility), five DNA tasks (spanning cell-type-specific enhancer activity, heat-shock-responsive promoters, and tissue-selective splicing), and three RNA design tasks (spanning translation efficiency, RNA folding, and toehold-switch function). Across these tasks, GRACE consistently achieves larger improvements in predicted target properties than discrete and continuous baselines while satisfying predicted auxiliary constraints. Together, these results establish GRACE as a general framework for machine-learning-guided biological sequence redesign under competing property constraints.

**Code and data availability:** The source code and data used in this study are available at https://github.com/Singh-Lab/GRACE.

## 1 Introduction

Recent advances in generative modeling, particularly diffusion and flow-matching approaches, have brought remarkable progress in **de novo** protein design, producing novel structures and functional binders [49, 5]. Yet generating a promising design does not guarantee that it will work experimentally: design campaigns often generate large candidate pools, filter them computationally, and screen the selected designs to find functional hits [39]. For many practical applications, such as improving the thermostability of an enzyme, enhancing binding affinity, or tuning the activity of a ribozyme, a natural sequence already exists that performs the desired function imperfectly, and **redesign** offers a more tractable path by constraining optimization to the neighborhood of a known functional sequence and leveraging evolutionary selection as an implicit prior on viability [36, 32]. Traditionally, this task has been tackled experimentally through rounds of mutagenesis and screening or selection, which can explore only a small fraction of the enormous combinatorial sequence space. Advances in high-throughput assays and machine learning have increasingly enabled predictive models of molecular properties (often called *oracles*) to guide sequence search towards promising variants before experimental testing [51, 41, 10].

This has given rise to *oracle-guided sequence optimization*, in which a learned model of sequence–property relationships guides search through a high-dimensional, discrete, and highly non-convex design space. A wide range of approaches have been developed, including reinforcement learning [3], black-box optimization [2], gradient-based continuous relaxations [29], Bayesian optimization in learned representations [43], evolutionary search [42, 4], generative approaches [28, 25], and adaptive sampling [12, 11]. These methods have enabled machine-learning-guided design across proteins, DNA, and RNA, including antibody engineering, cell-type-specific cis-regulatory design, and 5′ UTR optimization [31, 19, 44, 40, 13].

Most sequence-design problems, however, are not simply problems of maximizing a single score. A functional biological sequence typically already possesses many desirable properties that should not be sacrificed while improving another. A therapeutic protein may require improved activity while retaining stability and solubility; a regulatory element may need increased activity in one cellular context while preserving its activity in others; and an engineered RNA may need improved translation while preserving structural or regulatory properties. Large deviations in such auxiliary properties can render an otherwise high-scoring design biologically unusable [46, 47, 27, 53, 17].

We therefore focus on *reference-centered constrained sequence optimization*. Given a functional starting sequence, the goal is to improve one designated target property while requiring multiple auxiliary properties to remain within prescribed tolerances of their values in the starting sequence. Unlike multi-objective formulations that seek to improve several properties simultaneously, this setting treats auxiliary properties as quantities to preserve. Auxiliary property reference values and tolerances define biologically motivated two-sided bounds that allow targeted improvements while retaining useful properties of the starting sequence. This setting presents two computational challenges. First, each auxiliary property defines a nonlinear tolerance region, and the intersection of these regions can leave only a narrow set of admissible directions for improving the target. Second, the discrete nature of biological sequences means that an update appearing locally feasible in a continuous relaxation need not remain feasible after discrete sequence realization. Existing strategies for sequence redesign address aspects of these challenges. Gradient-based sequence optimizers can efficiently exploit differentiable oracle gradients to improve a target property, but provide no explicit mechanism for keeping auxiliary properties close to their reference values [28, 29]. Constraint-aware black-box, evolutionary, and sampling-based strategies can enforce feasibility through candidate filtering or acquisition [2, 43, 42], but do not exploit the local gradient geometry that differentiable oracles provide. Penalty-based approaches incorporate auxiliary constraints into the optimization objective, but allow sufficient target improvement to compensate for constraint violations [21, 22]. These limitations motivate the development of an approach that uses oracle gradients to improve the target property while limiting changes in auxiliary properties.

We introduce **GRACE** (**G**radient-**R**esolved **A**daptive **C**onstrained **E**xploration), a gradient-guided framework for reference-centered biological sequence redesign. GRACE combines null-space projection, which suppresses target-gradient directions that change auxiliary properties, with an augmented-Lagrangian correction that responds when constraints are violated. A straight-through estimator [8], applied to Gumbel– Softmax samples [**?**], connects predictor evaluations of discrete sequences to gradient updates in logit space. To our knowledge, GRACE is the first to combine these two forms of predictor-guided constraint control for discrete, reference-centered biological sequence redesign. We evaluate GRACE on 14 constrained sequence-design tasks spanning proteins, DNA, and RNA. These tasks cover diverse biological objectives, including activity, stability, solubility, aggregation, translation, RNA folding, enhancer specificity, splicing, and stimulus-responsive regulation, and involve substantially different sequence spaces, oracle architectures, and constraint structures. Across this broad benchmark, GRACE substantially outperforms gradient-based, penalty-based, evolutionary, black-box, and generative methods adapted to the same constrained optimization setting, achieving larger improvements in the predicted target property while satisfying all auxiliary constraints. Independent computational evaluations provide further support for the resulting designs beyond the predictors used during optimization. In sum, we formalize reference-centered constrained redesign as a general problem in biological sequence design, and the results establish GRACE as a broadly effective approach to improving a desired biological function while preserving other essential properties.

## 2 Problem Statement

We formulate reference-centered constrained sequence optimization as follows. For a natural sequence *x*_0_ ∈ {0, 1}^L×A^ of length *L* over an alphabet of size *A*, the goal is to find edits that improve one target property while preserving a set of auxiliary properties.

We assume access to a differentiable target oracle *f*_T_ (*x*), which scores the property to be improved, and constraint oracles 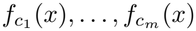, which measure auxiliary properties that should remain close to their values on the starting sequence *x*_0_. Formally, we seek to maximize the target property subject to auxiliary-property constraints:

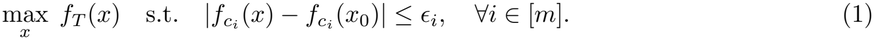

where *ϵ*_i_ is the allowed deviation for the *i*-th auxiliary property. Thus, a sequence is desirable only if it improves the target while remaining close to the wild-type auxiliary-property profile. We note that this framework naturally handles multiple target properties as well.

## 3 Method

### Overview of framework

Given datasets for the target and auxiliary properties, we first train machine learning models to predict these properties (Figure 1A). Alternatively, we can use pretrained models, provided we have access to the gradients needed for optimization. We then hold the predictors fixed and provide them, together with the reference sequence, to GRACE, which searches for a sequence that improves the predicted target property while keeping predicted auxiliary properties within specified tolerances of their reference values (Figure 1B). GRACE represents the sequence using continuous, trainable logits and samples discrete sequences for evaluation by the property predictors, using a straight-through estimator (STE) to obtain gradients with respect to the logits (Figure 1C). During the search, GRACE projects the target gradient to suppress components that change auxiliary properties to first order (Figure 1D) and uses an augmented Lagrangian mechanism to correct constraint violations (Figure 1E).

**Figure 1.**
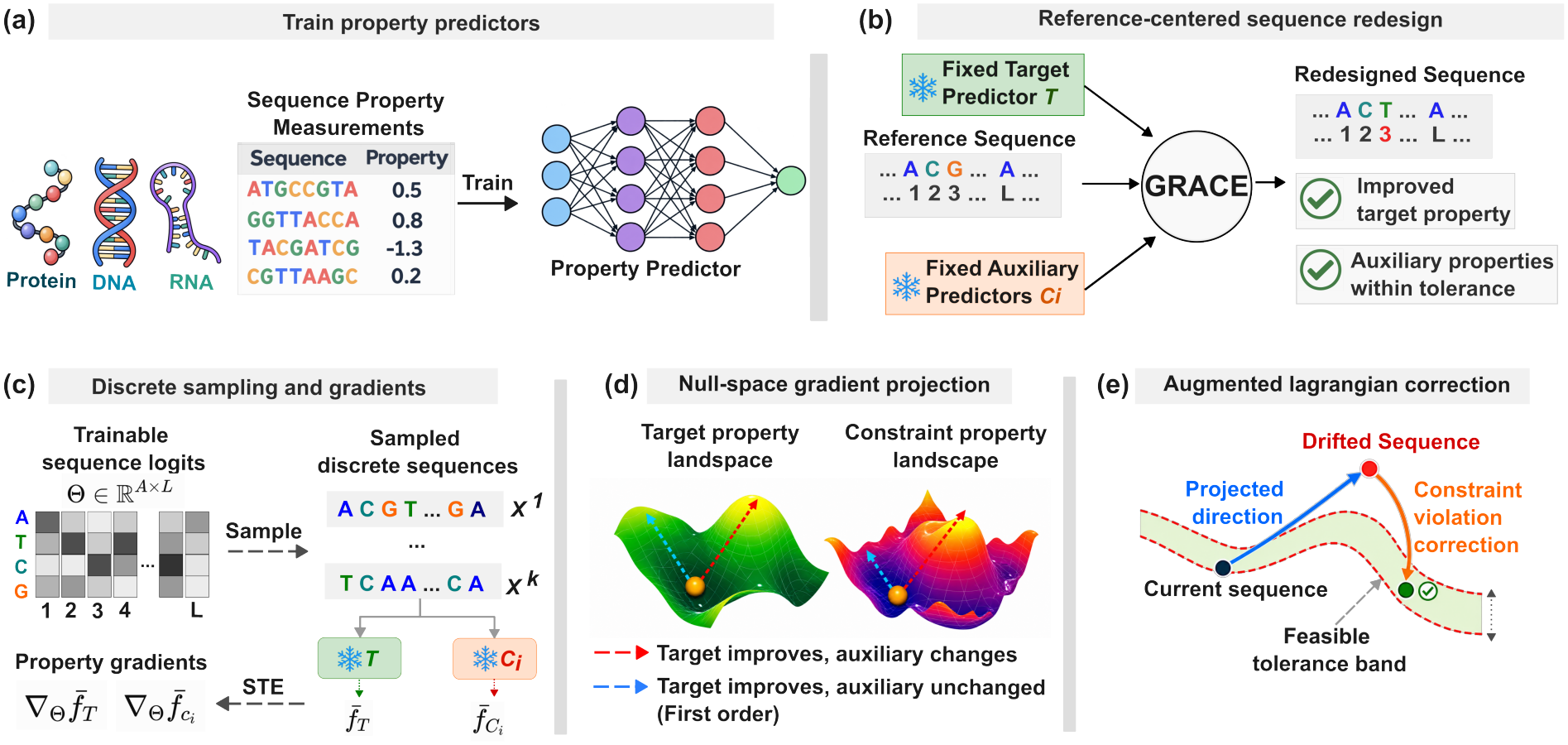
Overview of the GRACE framework for constrained biological sequence optimization. (a) Sequence–property measurements for proteins, DNA, and RNA are used to train property predictors. (b) Starting from a reference sequence, GRACE improves a target property while maintaining auxiliary properties within specified tolerances, using fixed predictors. (c) Trainable sequence logits generate discrete candidates, with a straight-through estimator (STE) enabling gradient propagation through sampling. (d) Null-space projection removes components of the target gradient that would cause first-order changes in auxiliary properties. (e) Augmented-Lagrangian correction addresses residual constraint violations arising from nonlinear drift, guiding optimization toward feasible sequences with improved target properties.

### 3.1 Gradient-Guided Optimization and Local Constraint Drift

We briefly review classical constrained optimization principles in the context of GRACE to motivate its design.

Suppose differentiable predictors evaluate a desired target property and auxiliary properties to be preserved. Their gradients provide directions for improving biological sequences, as used in existing gradient-based methods [29]. We represent a sequence by trainable logits Θ ∈ ℝ^A×L^, where *A* is the alphabet size and *L* the sequence length; each entry controls the preference for a sequence element at one position. Let 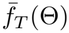 and 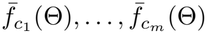 denote the target and auxiliary predictions as differentiable functions of these logits (Section 3.4). Their gradients describe local changes in the predicted properties. In particular, 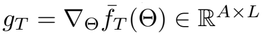 points toward local target improvement.

For an update Θ^+^ = Θ + *ηd*, where *η >* 0 is the step size, a first-order Taylor expansion gives

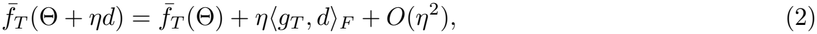

where ⟨·, ·⟩_F_ is the Frobenius inner product. Without constraints, *d* = *g*_T_ gives a first-order improvement of 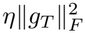, but may also change auxiliary properties (Fig. 1d, second panel). A standard approach in constrained optimization is to project the target gradient onto directions orthogonal to the relevant constraint gradients [37, 38]. For each selected constraint *i*, this requires 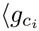, *d*⟩_F_ = 0, where 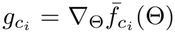. Suchdirections lie in the constraint Jacobian’s null space and preserve the selected properties to first order. Because auxiliary properties may vary within their tolerance bands, GRACE selects constraints for projection as they approach their boundaries. Sections 3.2 and 3.5 describe these steps.

First-order preservation, however, does not ensure that auxiliary properties remain within their bounds after a finite update. In nonlinear constrained optimization, locally feasible directions can drift away from curved constraint surfaces [34, 50] (Fig. 1d, third panel). A second-order Taylor expansion shows why:

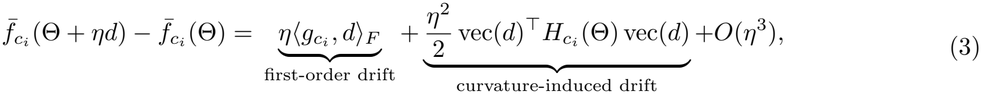

where 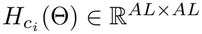 is the Hessian with respect to vectorized logits, assuming sufficient local smoothness. Even if the first-order term vanishes, curvature can produce a nonzero change and push a property outside its tolerance.

These observations motivate two complementary requirements: directions that improve the target while limiting first-order changes in selected auxiliary properties, and a mechanism that corrects residual violations from curvature and finite steps. The latter also matters when high-dimensional logits are used to realize discrete sequences, for which local approximations need not guarantee feasibility. GRACE obtains surrogate gradients through discrete evaluations using a straight-through estimator; the smooth analysis provides intuition, not a guarantee about changes in discrete predictions. Sections 3.3 and 3.4 describe violation correction and discrete sequence optimization, respectively.

### 3.2 Eliminating First-Order Drift by Null-Space Projection

We address the first-order drift in Eq. (3) by projecting the target gradient onto the constraint Jacobian’s null space [37, 38]. Here, logits and gradients are implicitly vectorized in ℝ^AL^. Let 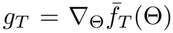 and 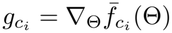. Stacking the constraint gradients gives

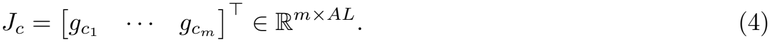

For a direction *d, J*_c_*d* collects the directional derivatives of the constrained properties; thus *J*_c_*d* = 0 preserves them to first order. When *J*_c_ has full row rank, the orthogonal projection of *g*_T_ onto Null(*J*_c_) is

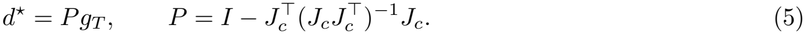

**Proposition 1 (Null-space projection eliminates first-order constraint drift)**. For the direction *d*^⋆^ defined in Eq. (5), *J*_c_*d*^⋆^ = 0. Consequently, assuming sufficient local smoothness, for every constraint *i*, 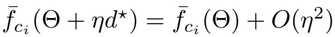, so the projected update preserves all constrained properties to first order. This follows by substituting the definition of *P* :

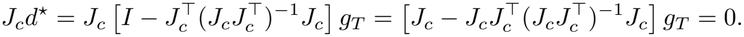

Geometrically, each constraint gradient points locally in a direction that changes the corresponding auxiliary property. Their joint null space therefore forms the local tangent subspace along which the constrained properties do not change to first order. GRACE follows the projection of the target gradient onto this subspace, retaining the closest direction to the target gradient that preserves the auxiliary properties to first order. Redundant or nearly collinear constraint gradients can make 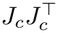 singular or ill-conditioned. We therefore use a damped projection with *δ* = 10^™6^ [48, 16]:

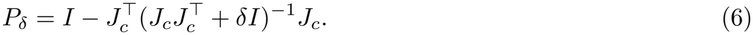

For full-row-rank *J*_c_, *P*_δ_ approaches the exact null-space projector as *δ* → 0. At finite *δ*, it suppresses but generally does not eliminate first-order constraint drift (Appendix 7.7); it also cannot eliminate the curvature-induced drift in Eq. (3). The next section addresses the resulting constraint violations.

### 3.3 Correcting Drift with an Augmented Penalty

Exact null-space projection eliminates first-order constraint drift, but it does not guarantee feasibility over finite update steps. Because the property predictors are nonlinear, curvature can cause an update along the projected direction to move outside the feasible region, even when the direction is locally constraint-preserving (Figure 1E). To address constraint violations, GRACE introduces a reactive correction that activates when a predicted auxiliary property exceeds its allowed tolerance. Let 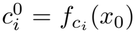 denote the value of the *i*th auxiliary property in the starting sequence. To measure how far a property has moved beyond its allowed range, we define its signed constraint residual as 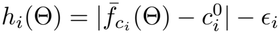, and its nonnegative violation as *v*_i_(Θ) = max(0, *h*_i_(Θ)). This violation is zero as long as the predicted property remains within its specified tolerance, and becomes positive only when that tolerance is exceeded. The absolute value accounts for deviations on either side of the reference value, while clipping at zero ensures that properties within their tolerance bands incur no penalty. To bring violated constraints back toward their allowed ranges, GRACE uses a modified augmented Lagrangian penalty that combines adaptive multiplier terms with quadratic penalties on constraint violations:

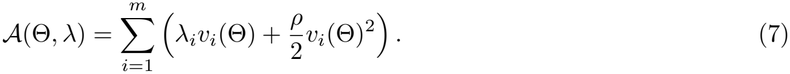

Here, *λ*_i_ ≥ 0 controls the linear penalty for the *i*th constraint, and *ρ >* 0 determines the strength of the quadratic penalty. The linear terms play the role of Lagrange-multiplier terms, while the quadratic terms augment them to strengthen the correction as violations grow. Unlike a standard inequality augmented Lagrangian written in terms of signed constraint residuals, this construction operates on nonnegative, clipped violations. The multipliers *λ*_i_ are updated across iterations, where positive constraint residuals increase them and negative residuals decrease them. The update scheme for the *λ*_i_ is given in Appendix 7.2.

The gradient of this penalty provides a corrective direction whenever a constraint is violated. GRACE combines this correction with the projected target gradient to obtain the final update:

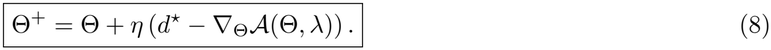

When all constraints are satisfied, the correction vanishes and optimization follows the projected target direction *d*^⋆^. When a violation occurs, the penalty gradient encourages a return toward the feasible region.

### 3.4 Gradient Optimization of Discrete Sequences

The preceding derivation operates on continuous logits Θ ∈ ℝ^A×L^, although biological sequences are discrete. We treat each column of Θ as trainable preferences for the nucleotides or amino acids at one position; a column-wise softmax defines categorical sampling probabilities. At each iteration, we draw *K* sequences. For sample *k*, Gumbel–Softmax produces a discrete one-hot sequence 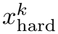 and a differentiable soft representation *P* ^k^ from the same perturbed logits [29]. Its temperature controls how closely *P* ^k^ approaches a one-hot representation. We combine them with a straight-through estimator (STE):

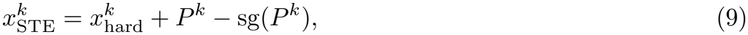

where sg preserves values but stops gradients. Thus, 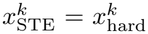 in the forward pass, so the predictors evaluate discrete sequences, while 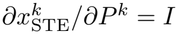 in the backward pass, so gradients flow through *P* ^k^ to Θ.

We average predictions across samples: 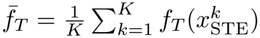 and 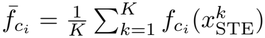. Backpropagation yields the surrogate gradients 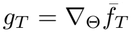 and 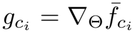 used to compute Θ^+^ = Θ + *ηd*_GRACE_. Accordingly, optimization constrains sample-averaged auxiliary predictions, rather than every sampled sequence individually. Further sampling details appear in Appendix 7.6.

### 3.5 Slack-Aware Projection

Projecting against constraints far from their tolerance boundaries can unnecessarily limit target improvement. GRACE therefore selects constraint *i* for projection when 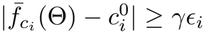, where *γ* ∈ [0, 1] determines how much of the tolerance band may be used before projection begins. Let *J*_active_ contain the gradients of the selected constraints. The projected direction is

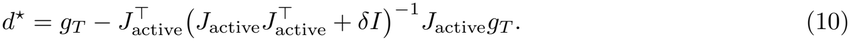

If none are active, *d*^⋆^ = *g*_T_. Setting *γ* = 0 selects all constraints; *γ* = 1 selects only those at or beyond their tolerance boundaries. Selection affects only projection. The full algorithm and hyperparameters appear in Appendix 7.

### 3.6 Experimental Setup

#### Benchmark tasks

We curate a benchmark of 14 constrained sequence optimization tasks spanning proteins, RNA, and DNA, with diverse functions, sequence lengths, and property predictors. Each task improves a target property while keeping auxiliary properties of a starting sequence within specified tolerances. We use trained or pretrained predictors alongside directly computed properties, all differentiable with respect to the sequence representation, and newly train 17 predictors for this benchmark. We summarize the tasks below; datasets, property calculations, and predictor training are detailed in Appendix 1.

For **proteins**, we consider six tasks. Two involve TEM-1 *β*-lactamase, a well-studied antibiotic-resistance enzyme, where we improve either enzymatic activity or solubility while preserving the other property and folding stability, using predictors trained on experimental mutational data. We also optimize catalytic turnover (*k*_cat_) using enzyme sequences with measured turnover numbers [24], while preserving hydrophobicity. For aggregation, we use short peptides experimentally screened for amyloid nucleation in the CANYA dataset [45], aiming to reduce nucleation while preserving net charge. For solubility, we use proteins from the *E. coli* proteome with experimentally measured solubility [33], aiming to improve solubility without altering net charge. Finally, we consider thousands of computationally designed mini-proteins experimentally tested for folding stability by Rocklin et al. [35], aiming to improve stability while preserving net charge.

For **RNA**, we consider three tasks. First, we use synthetic human 5′UTRs with experimentally measured ribosome loading [40], aiming to increase translation efficiency while preserving folding energy. Second, we optimize synthetic toehold switches, which regulate expression in response to a trigger RNA, using measured ON- and OFF-state fluorescence [1]. The goal is to increase ON-state expression while preserving OFF-state leakage. Third, we optimize RNA folding stability using a surrogate model trained on folding free energies computed with ViennaRNA [30], while preserving GC content. For **DNA**, we consider five tasks. Three involve human regulatory sequences with experimentally measured enhancer activities in K562, HepG2, and SK-N-SH cells [19]. We increase activity in one cell type while preserving activity in the other two. We also consider human promoters assayed under normal and heat-shock conditions [7], aiming to increase heat-shock inducibility while preserving basal expression. Finally, using human splice sites and Pangolin [52], a predictor of tissue-specific splice-site usage, we seek to increase splice-site usage in liver while preserving usage in heart, brain, and testis. For most protein, RNA, and enhancer tasks, we select 30 starting sequences with room for target improvement (15th–35th percentile of the candidate pool) and intermediate constraint values (20th–60th percentile), enforcing a minimum Hamming distance between selections. TEM-1 redesigns start from the wild-type enzyme. For heat-shock regulation, we select 30 held-out promoters with measurable basal activity and low predicted inducibility from 80 eligible promoters; for splicing, we use 48 held-out human splice sites with intermediate liver usage. We run GRACE twice per start with different seeds, except for TEM-1 (eight seeds), heat-shock regulation (five seeds), and splicing (one run per site). For each run, we record the largest predicted target improvement over the start among individually evaluated discrete candidates predicted to satisfy all auxiliary constraints. We note that, experimental validation is needed to establish whether these gains and constraint satisfaction hold in practice. We collapse runs from each start to their median, then report the median and interquartile range across starting sequences. The benchmark, including processed datasets, trained predictors, starting sequences, and evaluation protocols, is publicly released. Further details appear in Appendix 1.

#### Baseline comparator methods

We compare GRACE against established sequence optimization methods spanning four optimization paradigms. Since these methods were not originally designed to improve a target property while preserving multiple auxiliary properties of a reference sequence within two-sided tolerances, we adapt their standard implementations to accommodate our constraints. For direct gradient optimization, we use **Fast SeqProp** [29], which maximizes the target property; we filter its sequences for feasibility. For evolutionary and stochastic search, we adapt **AdaLead** [42], **directed evolution** [4], **simulated annealing** [23], and **random search** [9]. For all four, candidates are checked against the auxiliary constraints at each search step and the best feasible variant is retained. For **constrained Bayesian optimization** [18, 6], we fit Gaussian process surrogates to the target and auxiliary scores and use a feasibility-aware acquisition function balancing expected improvement against the probability of satisfying all constraints, scored over discrete pools of mutants of the reference and promising candidates. For **generative approaches**, we train a task-specific masked-denoising convolutional network on each sequence library. Following an iterative inpainting strategy [14], we mask 5–30% of positions and refill them over eight steps, guiding the denoiser’s output probabilities with the gradient of the target score penalized by two-sided constraint violations. We compare **guided discrete diffusion**, which applies this guidance directly, with **prior-preserving guided diffusion**, which adds a stronger likelihood term to keep sampling closer to the learned sequence prior [20]. Generated sequences are filtered for feasibility. For the three enhancer tasks, we also compare with **Ctrl-DNA** [15], constrained reinforcement learning whose original objective promotes target-cell specificity rather than preserving off-target activities within two-sided tolerances. Because it is sample-inefficient, we report it at the matched budget (**Ctrl-DNA**) and at 5× that budget (**Ctrl-DNA (5**× **budget)**). Budgets are measured in property-predictor calls. All methods use the same starting sequences and frozen property oracles under identical feasibility criteria; detailed adaptations, predictor-call counts, and hyperparameters are in Appendix 2 and 6.

## 4 Results

### 4.1 GRACE improves protein properties while preserving auxiliary constraints

We evaluate GRACE on six protein optimization tasks spanning enzymatic activity, solubility, catalytic turnover, amyloid nucleation, and protein stability (Fig. 2). The tasks use predictors trained on experimental measurements as well as exact sequence-derived properties (e.g., net charge and hydrophobicity). While the two TEM-1 tasks use the same starting sequence, with eight runs per task using different random seeds, the remaining four tasks use 30 distinct starting sequences with two stochastic runs per starting sequence. For each task, we report predicted target property values and assess whether the auxiliary properties satisfy their specified constraints according to the corresponding predictors.

**Figure 2.**
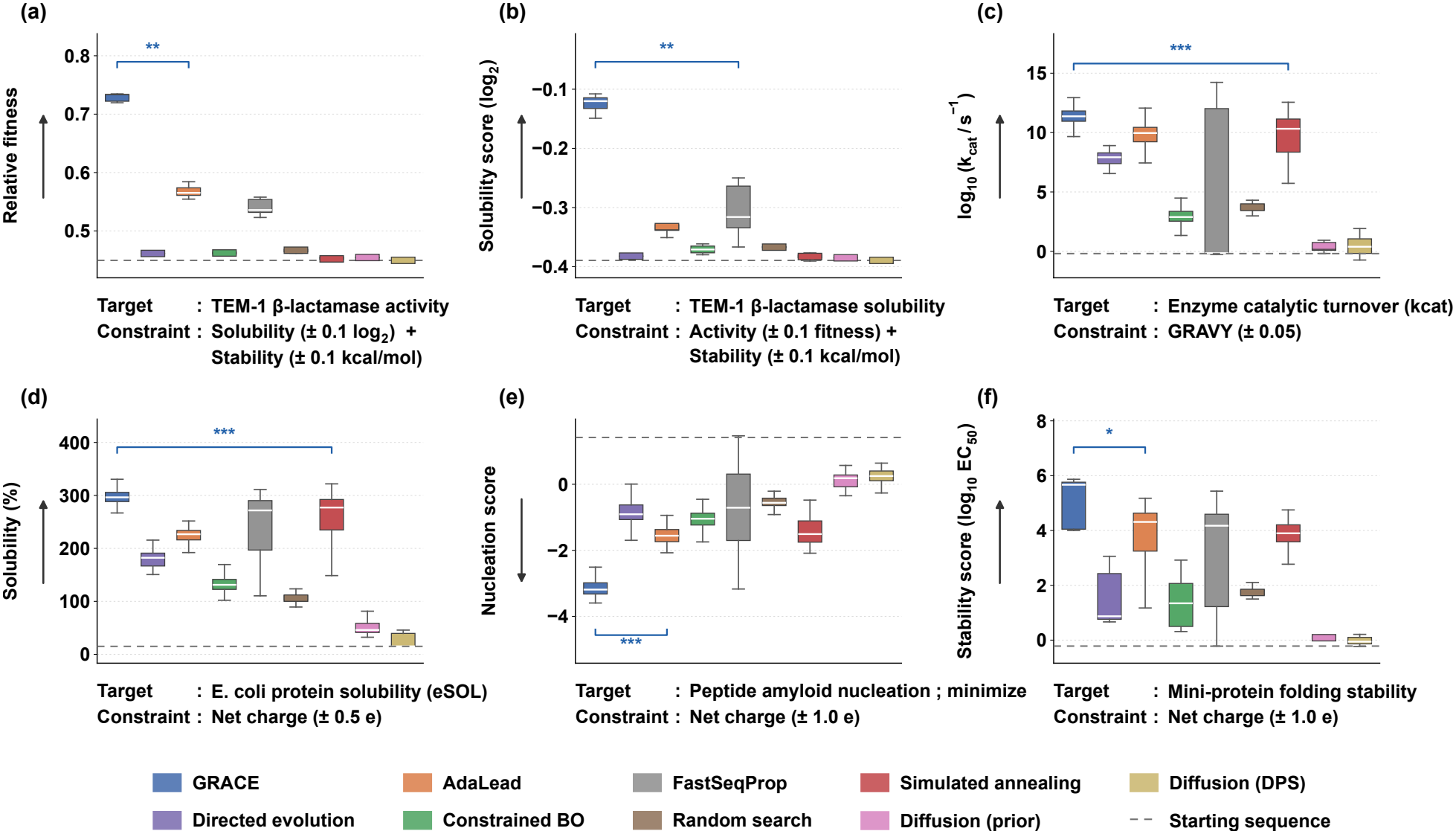
GRACE improves target properties while preserving constraints across diverse protein-design tasks. **(a)** TEM-1 activity, with solubility and stability preserved (*±*0.1). **(b)** TEM-1 solubility, with activity and stability preserved (*±*0.1). **(c)** Catalytic turnover (log_10_ *k*_cat_), with hydrophobicity (GRAVY) preserved (*±*0.05); these windows are all 40 aa, so this equals the *±*2.0 summed Kyte–Doolittle tolerance used in the supplement. **(d)** Protein solubility, with net charge preserved (*±*0.5). **(e)** Amyloid nucleation minimization, with net charge preserved (*±*1.0). **(f)** Folding-stability maximization, with net charge preserved (*±*1.0). Boxplots show the predicted target score of the best feasible sequence from each independent run (one score per run). The TEM-1 tasks use the wild-type reference with eight seeds (*n* = 8); other tasks use 30 starts with two seeds each (*n* = 60), except guided diffusion, which uses a subset of starts (*n* = 12–15). Feasibility requires every auxiliary property to remain within its stated tolerance of the reference value. Arrows indicate whether the target is maximized (*↑*) or minimized (*↓*); dashed lines mark the mean predicted target score of the starting sequences. Paired Wilcoxon signed-rank tests compare GRACE with the strongest baseline for each task, pairing by start after collapsing seeds to their per-start median (^*\**^*p <* 0.05, ^*\*\**^*p <* 0.01, ^*\*\*\**^*p <* 0.001; n.s., not significant).

**Figure 3.**
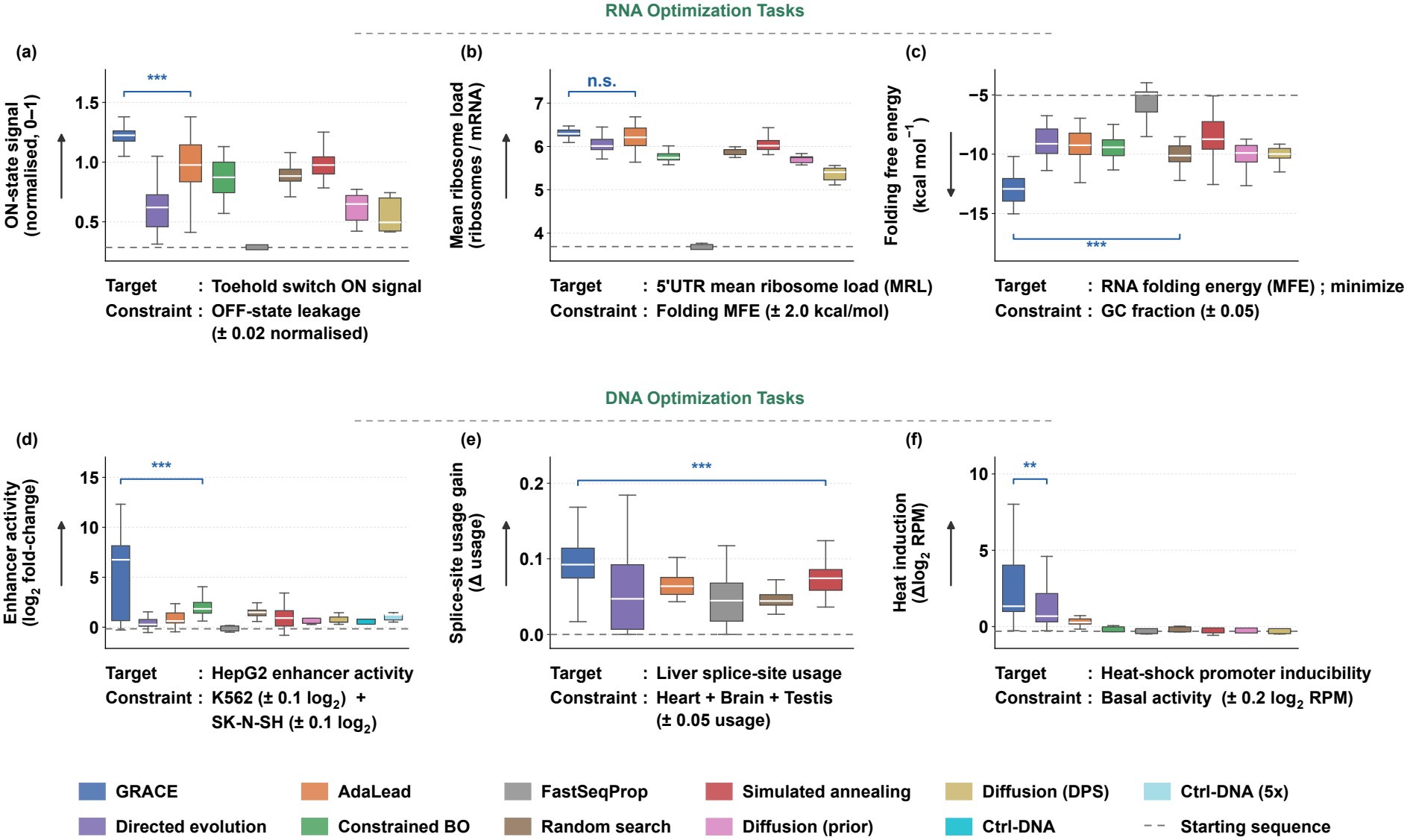
GRACE improves nucleotide-sequence properties while preserving constraints across RNA and DNA design tasks. The top row shows RNA tasks; the bottom row shows DNA tasks. **(a)** Toehold-switch ON-state expression, preserving OFF-state leakage (*±*0.02). **(b)** 5^*′*^UTR mean ribosome loading (MRL), preserving folding free energy (*±*2.0 kcal/mol). **(c)** RNA folding free-energy minimization, preserving GC fraction (*±*0.05). **(d)** HepG2 enhancer activity, preserving K562 and SK-N-SH activity (each within *±*0.1). **(e)** Liver-specific splice-site usage, preserving heart, brain, and testis usage (each within *±*0.05) and allowing at most 30 substitutions. **(f)** Heat-shock inducibility, preserving basal promoter activity (*±*0.2) and allowing at most 10 substitutions. Boxplots show the predicted target score of the best feasible sequence per independent run. Panels (a)–(d) use 30 starts and two seeds (*n* = 60), with *n* = 50–59 for constrained Bayesian optimization, *n* = 15 for guided diffusion, and *n* = 5 for Ctrl-DNA. Panel (e) uses 48 held-out splice sites with one run each; panel (f) uses 30 promoters and reports the best of five restarts per promoter. Feasibility requires every auxiliary property to remain within its stated tolerance of the reference value. Ctrl-DNA appears only in (d), as it is specific to regulatory DNA; guided diffusion is omitted from (e) because this task lacks the sequence library needed to train a task-specific denoiser. Arrows denote maximization (*↑*) or minimization (*↓*); dashed lines show the mean predicted starting score. Paired Wilcoxon signed-rank tests compare GRACE with each task’s strongest baseline, pairing starts after taking the median across seeds (^*\**^*p <* 0.05, ^*\*\**^*p <* 0.01, ^*\*\*\**^*p <* 0.001; n.s., not significant).

On all six tasks, GRACE achieves the highest median predicted feasible improvement as compared to other baseline methods, with statistically significant gains over the next-best method on each task (Wilcoxon signed-rank test). Relative to the strongest baseline for each task, GRACE’s improvements are 13%–265% larger. The largest advantages occur on predicted TEM-1 activity and solubility (141% and 265%) as well as amyloid nucleation (54%), while smaller but consistent gains are observed for protein solubility (13%) and catalytic turnover (14%). AdaLead is the strongest baseline on four of the six tasks and Fast SeqProp on the remaining two, with no method approaching GRACE on either TEM-1 task.

GRACE also generates constraint-satisfying candidates throughout optimization; this is a desirable property as each run proposes *multiple* sequences that improve the predicted target property while satisfying the auxiliary constraints. Across all six tasks, 26.0–83.9% of GRACE’s target-improving candidates satisfy the constraints, compared with only 1.2–6.2% for Fast SeqProp, which aims to find sequences that simply improve the target property (Appendix Table 16). Additionally, across all tasks, GRACE retains a median of 95.2–100% of its best discovered target improvement when restricting to sequences that satisfy the auxiliary constraints (Appendix Table 17). Moreover, among candidates that do not satisfy the auxiliary properties, GRACE’s violations are smaller than those of every evolutionary, stochastic, and gradient baseline on all six tasks; constrained Bayesian optimization and guided diffusion are comparable, consistent with the much smaller edits they propose (Appendix Table 18). The extent of sequence change varies across tasks, with GRACE finding feasible improvements at different levels of identity to the starting sequence (Appendix 3.5). For example, on TEM-1 activity (Fig. 4a), GRACE increases predicted activity from 0.449 to 0.734 by introducing 97 substitutions across the 285-residue protein. Despite these changes, all eight annotated catalytic and substrate-binding residues remain unchanged, although no explicit constraints were imposed to keep them unmutated. In contrast, the AdaLead design reaches an activity score of 0.584 but mutates the active-site residue N170. This example illustrates how GRACE can make extensive sequence changes while retaining key functional positions.

**Figure 4.**
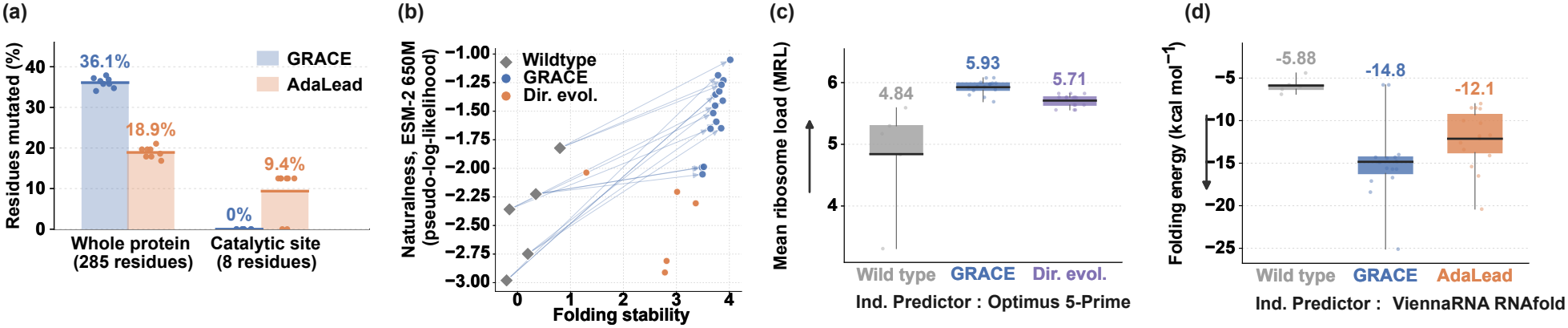
Biological characterization and independent computational evaluation of GRACE designs. These analyses use functional-site annotations or predictors excluded from optimization. **(a)** Fraction of TEM-1 residues mutated by GRACE and AdaLead, shown for the full protein and its eight annotated catalytic and substrate-binding residues. GRACE makes more substitutions overall while preserving all eight functional residues. Dots show individual runs; bars show means. **(b)** Mini-protein folding stability versus sequence naturalness, measured by ESM2-650M pseudo-log-likelihood. Lines connect starting sequences to GRACE designs; directed-evolution designs are shown for comparison. **(c)** Independent Optimus 5-Prime predictions of 5^*′*^UTR mean ribosome loading for wild-type, GRACE, and directed-evolution sequences. **(d)** Independent ViennaRNA RNAfold predictions of folding free energy for wild-type, GRACE, and AdaLead sequences; more negative values indicate more stable predicted RNA structures.

### 4.2 GRACE generalizes across diverse RNA and DNA tasks

We evaluate GRACE on three RNA tasks and three DNA tasks (Fig. 3), where improving one function can affect closely related properties. GRACE achieves the highest median feasible improvement across all six tasks, with a 2%–117% median paired relative advantage over the strongest competing baseline. The largest gains occur in heat-shock promoters (117%), HepG2 enhancers (116%), and RNA folding (57%). GRACE finds significantly better predicted target properties for five of six tasks, with 5′UTR translation being the exception (*p* = 0.05, Wilcoxon signed-rank test). Additional results for K562 and SK-N-SH enhancer optimization demonstrate similar advantages (Appendix Fig. 1a). GRACE achieves these improvements while preserving auxiliary properties, as illustrated by a therapeutic mRNA design example. An open 5′UTR can facilitate translation initiation, but openness alone does not ensure high ribosome loading [53, 13]. Starting from a 50-nt 5′UTR with an RNAfold-predicted minimum free energy of 0 kcal/mol and predicted mean ribosome loading of 0.612, GRACE introduces 15 substitutions that increase predicted ribosome loading to 6.16 while maintaining the minimum free energy at 0 kcal/mol (Appendix 9b). Similarly, GRACE improves DNA regulatory activity while preserving activity in non-target cell types, splice-site usage in other tissues, or basal promoter activity.

Trajectory analysis further demonstrates GRACE’s ability to control constraint violations. Across the eight RNA/DNA tasks, GRACE retains 86.2–100% of its best discovered improvement on seven tasks, with RNA folding being the exception (55.3%; Appendix Table 17). Among the evolutionary and stochastic search baselines-AdaLead, directed evolution, and simulated annealing - GRACE achieves the lowest violation severity on five tasks, including all three RNA tasks, splicing, and heat-shock promoters (Appendix Table 18). On the toehold task, for example, its mean violation severity is only 1.73 tolerance units, compared with 22.08 for AdaLead and 34.24 for FastSeqProp. On enhancer tasks, simulated annealing achieves higher trajectory feasibility and smaller violations, but produces substantially lower feasible improvements than GRACE (Appendix Tables 16 and 18). Importantly, these results do not arise from restricting optimization to small sequence changes. GRACE’s returned RNA and enhancer designs retain only 50.0–68.5% identity to their starting sequences. On enhancer tasks, GRACE also produces more diverse designs than AdaLead, with 71.5–76.5% versus 89.0–91.0% pairwise identity (Appendix Table 19). Moreover, GRACE requires only 7–27 s per run on CPU-based tasks, comparable to inexpensive search baselines and more than two orders of magnitude faster than constrained Bayesian optimization on the three RNA tasks, where both were run (Appendix Fig. 3). Together, these results demonstrate

### 4.3 Independent evaluations support the biological validity of GRACE designs

To assess whether GRACE’s improvements extend beyond the oracles used during optimization, we evaluate its designs with independent predictors, structural models, and biological annotations (Fig. 4). These evaluations examine whether designs retain structural and functional features and score favorably under alternative methods. For TEM-1, AlphaFold3 predictions support preservation of the overall fold (Appendix 9), while sequence analysis shows that all eight annotated functional residues remain unchanged, despite not being explicitly constrained (Fig. 4a). For mini-proteins from the Rocklin dataset [35], we train an independent stability predictor with a transformer backbone, rather than the CNN backbone used during optimization (Appendix 1). Under this predictor, GRACE designs show favorable predicted stability and higher ESM2-650M pseudo-log-likelihoods [26] than directed evolution, the strongest baseline (Fig. 4b). For 5′UTR design, the independent translation predictor Optimus 5-Prime [13] assigns GRACE designs higher mean ribosome loading than the strongest baseline (5.93 versus 5.71; Fig. 4c). For RNA folding designs, the physics-based ViennaRNA RNAfold model [30] predicts lower folding free energy than the strongest baseline (™14.8 versus ™12.1 kcal/mol; Fig. 4d). Together, these computational evaluations support GRACE’s improvements beyond the optimization oracles, although experimental validation is needed to establish their biological effects.

### 4.4 Ablation studies and tolerance robustness

Ablation studies confirm that combining null-space projection with augmented-Lagrangian (ALM) correction improves both optimization performance and constraint satisfaction. Across 13 tasks, full GRACE achieves higher feasible target improvement than projection-only on 11 tasks and ALM-only on eight tasks, tying ALM-only on three others (Appendix Table 21). The benefit is particularly striking for enhancer optimization, where GRACE achieves 20–194× greater feasible improvement than projection-only and consistently outperforms ALM-only. Beyond target improvement, GRACE achieves higher trajectory feasibility and lower constraint violations than projection-only on all 13 tasks, and higher feasibility than ALM-only on 11 tasks. These results establish that the two components work together to achieve stronger target optimization while maintaining tighter constraint control. This advantage persists across constraint tolerances. Across the nine tasks in the sweep and five tolerance settings (0.5–2×), GRACE outperforms projection-only in 40 of 45 and ALM-only in 32 of 45 comparisons of feasible target improvement (Appendix Table 20). Moreover, GRACE surpasses the strongest non-ablation search baseline on eight of those nine tasks at every tested tolerance, 5′UTR translation being the exception. Together, these results demonstrate that GRACE’s performance gains arise from the combined optimization strategy rather than a favorable choice of constraint tolerance.

## 5 Discussion

GRACE addresses a common problem in biological sequence redesign: improving one property while retaining others that make the starting sequence useful. Its components play complementary roles. Null-space projection limits local changes in selected auxiliary properties, while the adaptive penalty responds to violations arising from finite steps or discrete sampling. Ablation and tolerance analyses show that combining them yields stronger feasible improvements than either alone across varied constraint settings. Across protein, RNA, and DNA tasks spanning different functions, predictors, and constraint structures, GRACE achieves larger predicted improvements while satisfying auxiliary constraints. Evaluations using predictors and annotations excluded from optimization provide further support for the designs. Experimental testing is still needed to determine whether the predicted gains and preserved properties hold in biological systems. GRACE also depends on accurate predictors with informative gradients. Although it produced diverse enhancer designs in our experiments, diversity is neither an explicit objective nor guaranteed: independent runs may converge on similar sequences. An explicit diversity objective could help generate distinct candidates. The framework could also extend to multi-objective de novo design: once a property reaches a sufficient threshold, treating it as a one-sided constraint would allow search to focus on unmet objectives while preserving that gain. Together, these results position GRACE as a broadly applicable framework for improving biological sequences under competing property requirements.

