## Supplementary Information for "GRACE: Discrete Biological Sequence Optimization with Gradient-Resolved Adaptive Constrained Exploration"

### Supplementary Material for
### GRACE: Gradient-Resolved Adaptive Constrained Exploration for Biological Sequence Redesign

**4 Robustness to Constraint Tolerances** **16**

**5 Component Ablation** **17**

**6 Fairness of Baseline Comparison** **18**

**7 GRACE Implementation Details** **19**

**8 Statistical Analysis** **23**

**9 Additional Results** **24**

**10 Computational Resources and Runtime** **25**

**11 Reproducibility and Public Release** **25**

### 1 Benchmark Tasks, Datasets, and Property Oracles

We evaluate GRACE on 14 constrained sequence-redesign tasks spanning proteins, RNA, and DNA. Each task specifies one target property to optimize and one or more auxiliary properties that must remain within prescribed tolerances of their values in the starting sequence. Property oracles are either learned predictors or exact sequence-derived functions. All learned predictors are frozen throughout optimization.

The following task profiles describe the biological motivation, optimization objective, constraint definitions, predictor training and evaluation, and starting-sequence selection. Target percentiles for minimization tasks are reported after orienting the objective so that higher values indicate better performance.

#### 1.1 Protein Tasks

Table 1: **TEM-1 beta-lactamase activity**

| | |
|---|---|
| **Task description** | **Motivation:** TEM-1 beta-lactamase enables bacterial resistance to ampicillin. The aim is to improve resistance-associated activity without sacrificing solubility or folding stability, since an active enzyme must remain soluble and properly folded.<br>**Target:** Maximize predicted ampicillin-resistance fitness (relative, dimensionless).<br>**Constraints:** Preserve solubility within ±0.1 $\log_2$ enrichment units and folding stability within ±0.1 kcal/mol of their starting values; at most 171 substitutions. |
| **Trained predictors** | **Dataset:** Activity: 5,198 TEM-1 single-mutant measurements [7]. Solubility: 4,130 yeast-display variants [12]. Stability: 471,684 training variants spanning 432 protein domains from Tsuboyama et al. (2023).<br>**Training splits:** Activity and solubility use five-fold position-based holdouts. Stability uses a domain-level split with 47 entirely unseen test domains.<br>**Model:** Separate fine-tuned ESM-2 protein language models, frozen during optimization.<br>**Performance:** Mean cross-fold Spearman $\rho = 0.494 \pm 0.077$ for activity and $\rho = 0.519 \pm 0.059$ for solubility; stability Pearson $r = 0.685$ on unseen domains. |
| **Starting sequences** | **Selection rationale:** One fixed TEM-1 reference provides a common, biologically recognizable starting point. It was not selected using the percentile-based starting-sequence procedure.<br>**Number and values:** One 285-aa reference. Frozen-oracle predictions are 0.450 for activity, −0.389 for solubility, and 1.922 for stability. These are predictions, not experimental measurements of this exact reference.<br>**Sequence diversity:** Pairwise Hamming distance is undefined for one start. |

Table 2: **TEM-1 beta-lactamase solubility**

| | |
|---|---|
| **Task description** | **Motivation:** Improving TEM-1 solubility can increase its functional availability and expression. However, solubility improvements should not compromise resistance-associated activity or folding stability.<br>**Target:** Maximize predicted yeast-display solubility ($\log_2$ enrichment).<br>**Constraints:** Preserve relative ampicillin-resistance fitness within ±0.1 and folding stability within ±0.1 kcal/mol of the start; at most 171 substitutions. |
| **Trained predictors** | **Dataset:** Solubility: 4,130 TEM-1 yeast-display variants [12]. Activity: 5,198 single mutants [7]. Stability: 471,684 variants from 432 protein domains, Tsuboyama et al. (2023).<br>**Training splits:** Five-fold position-based holdout for activity and solubility; domain-level holdout for stability, including 47 unseen test domains.<br>**Model:** Separate fine-tuned ESM-2 models for solubility, activity, and stability.<br>**Performance:** Solubility Spearman $\rho = 0.519 \pm 0.059$; activity $\rho = 0.494 \pm 0.077$; stability Pearson $r = 0.685$. |
| **Starting sequences** | **Selection rationale:** The same fixed TEM-1 reference is used as in the activity task, so only the target and constraint roles change. It was not percentile-selected.<br>**Number and values:** One 285-aa reference. Predicted starting solubility is −0.389, activity is 0.450, and stability is 1.922.<br>**Sequence diversity:** Pairwise Hamming distance is undefined for one start. |

Table 3: **Catalytic turnover**

| | |
|---|---|
| **Task description** | **Motivation:** Catalytic turnover, $k_{cat}$, measures how rapidly an enzyme converts substrate into product. Improving turnover is useful in enzyme engineering, but excessive changes in hydrophobicity may affect expression and biophysical behavior.<br>**Target:** Maximize predicted $\log_{10}(k_{cat}/s^{-1})$.<br>**Constraint:** Preserve summed Kyte–Doolittle hydrophobicity within ±2.0 scale units of the start. Every sequence in this task is 40 aa, so this is the same constraint as ±0.05 GRAVY (the length-averaged Kyte–Doolittle value, 2.0/40), which is how it is stated in Fig. **??**c; the optimiser applies the summed form. |
| **Trained predictors** | **Dataset:** 18,304 40-aa sequence windows from a curated enzyme-turnover collection derived from DLKcat [13]. Hydrophobicity is calculated directly from amino-acid identities.<br>**Training split:** Random 90/10 training/validation split (seed 0); no separate local test partition.<br>**Model:** One-dimensional CNN for catalytic turnover; exact hydrophobicity calculation.<br>**Performance:** Validation Spearman $\rho = 0.270$ for turnover; the constraint is exact. |
| **Starting sequences** | **Selection rationale:** Select sequences with low-to-moderate predicted turnover and intermediate hydrophobicity, leaving room for improvement while avoiding extreme constraint values. Favor distinct starts.<br>**Number and distribution:** Thirty 40-aa windows. Predicted turnover spans the 23rd–27th percentiles and hydrophobicity the 20th–59th percentiles of a 6,000-sequence candidate pool.<br>**Sequence diversity:** Median pairwise Hamming distance of 38 out of 40 positions across 435 pairs. |

Table 4: **Protein solubility**

| | |
|---|---|
| **Task description** | **Motivation:** Protein solubility affects recombinant production, formulation, and functional availability. Increasing charge can improve apparent solubility, so the charge constraint forces optimization to exploit other sequence determinants.<br>**Target:** Maximize predicted eSOL solubility score (dataset score units).<br>**Constraint:** Preserve computed net charge within ±0.5 elementary-charge units of the start. |
| **Trained predictors** | **Dataset:** 32,943 protein windows derived from the *E. coli* eSOL collection [16]. Net charge is computed directly.<br>**Training split:** Random 90/10 training/validation split (seed 0); no separate local test partition.<br>**Model:** One-dimensional CNN for solubility; exact amino-acid charge calculation.<br>**Performance:** Validation Spearman $\rho = 0.457$ for solubility; charge is exact. |
| **Starting sequences** | **Selection rationale:** Choose low-to-moderate predicted solubility with intermediate charge, avoiding constraint extremes and retaining improvement headroom.<br>**Number and distribution:** Thirty 40-aa windows. Target scores span the 23rd–28th percentiles and net charge the 17th–49th percentiles of the candidate pool.<br>**Sequence diversity:** Median pairwise Hamming distance of 37 out of 40 positions across 435 pairs. |

Table 5: **Amyloid nucleation**

| | |
|---|---|
| **Task description** | **Motivation:** Amyloid-forming peptides can nucleate aggregates associated with protein-misfolding diseases and biotechnology-related instability. Simply increasing charge can suppress aggregation, so charge is preserved to encourage more specific sequence changes.<br>**Target:** Minimize predicted CANYA amyloid nucleation score (assay score units).<br>**Constraint:** Preserve computed net charge within ±1.0 elementary-charge units of the start. |
| **Trained predictors** | **Dataset:** 42,235 random 20-aa peptides from the CANYA amyloid-nucleation assay [21]. Charge requires no trained predictor.<br>**Training split:** Random 90/10 training/validation split (seed 0); no separate local test partition.<br>**Model:** One-dimensional CNN trained on nucleation labels; exact charge calculation.<br>**Performance:** Validation Spearman $\rho = 0.354$ for nucleation; charge is exact. |
| **Starting sequences** | **Selection rationale:** Select peptides with room to reduce nucleation and intermediate charge. Diverse starts explore distinct sequence neighborhoods.<br>**Number and distribution:** Thirty 20-aa peptides. Target ranks span the 23rd–26th percentiles when lower nucleation is counted as better; charge spans the 18th–50th percentiles of the candidate pool.<br>**Sequence diversity:** Median pairwise Hamming distance of 19 out of 20 positions across 435 pairs. |

Table 6: **Designed mini-protein stability**

| | |
|---|---|
| **Task description** | **Motivation:** Designed mini-proteins provide compact scaffolds for protein engineering, but maintaining a stable fold is sensitive to sequence changes. The charge constraint limits improvements arising from large changes in electrostatic character.<br>**Target:** Maximize predicted proteolysis-derived stability score (assay score units).<br>**Constraint:** Preserve computed net charge within ±1.0 elementary-charge units of the start. |
| **Trained predictors** | **Dataset:** 48,399 designed 43-aa proteins from Rocklin et al. [18], distributed through the TAPE stability benchmark [17].<br>**Training split:** Random 90/10 training/validation split (seed 0); no separate local test partition.<br>**Model:** One-dimensional CNN for stability; exact charge calculation.<br>**Performance:** Validation Spearman $\rho = 0.755$; charge is exact. |
| **Starting sequences** | **Selection rationale:** The same percentile rule as the other CNN tasks: low-to-moderate predicted stability for headroom, with net charge away from the extremes.<br>**Number and distribution:** Thirty 43-aa proteins. Under the frozen predictor, target ranks span the 24th–27th percentiles and charge the 15th–59th percentiles of the 6,000-sequence candidate pool.<br>**Sequence diversity:** Median pairwise Hamming distance of 38 out of 43 positions across 435 pairs. |

#### 1.2 RNA Tasks

Table 7: **Toehold switch**

| | |
|---|---|
| **Task description** | **Motivation:** A toehold switch should produce a strong signal when its cognate trigger is present while remaining inactive without the trigger. Increasing ON-state output at the expense of higher OFF-state leakage would produce a poor switch.<br>**Target:** Maximize normalized ON-state output (dimensionless, 0–1).<br>**Constraint:** Maintain normalized OFF-state leakage within ±0.02 of its starting value. |
| **Trained predictors** | **Dataset:** 60,750 ON-labelled and 78,455 OFF-labelled switches from the experimental library of Angenent-Mari et al. [1].<br>**Training split:** Separate random 90/10 training/validation splits for ON and OFF measurements (seed 0).<br>**Model:** Separate one-dimensional CNN predictors for ON output and OFF leakage.<br>**Performance:** Validation Spearman $\rho = 0.660$ for ON output and $\rho = 0.524$ for OFF leakage. |
| **Starting sequences** | **Selection rationale:** Choose low-to-moderate ON output to allow improvement while keeping OFF leakage away from extreme values. Favor distinct sequences.<br>**Number and distribution:** Thirty 30-nt switches. Predicted ON scores span the 22nd–27th percentiles and OFF scores the 21st–59th percentiles of the candidate pool.<br>**Sequence diversity:** Median pairwise Hamming distance of 21 out of 30 positions across 435 pairs. |

Table 8: **5′UTR translation**

| | |
|---|---|
| **Task description** | **Motivation:** The 5′UTR regulates translation initiation and ribosome loading, making it important for synthetic mRNAs and mRNA therapeutics. The goal is to increase translation without simply removing RNA structure.<br>**Target:** Maximize predicted mean ribosome load (MRL, ribosomes per transcript).<br>**Constraint:** Preserve predicted minimum folding free energy within ±2.0 kcal/mol of the start. |
| **Trained predictors** | **Dataset:** 30,000 50-nt UTRs with MRL labels from the Optimus 5-Prime setting [19] and MFE labels computed with ViennaRNA [15].<br>**Training split:** Random 90/10 training/validation split (seed 0); no separate local test partition.<br>**Model:** Separate one-dimensional CNN predictors for MRL and folding energy.<br>**Performance:** Validation Spearman $\rho = 0.838$ for MRL and $\rho = 0.747$ for MFE. |
| **Starting sequences** | **Selection rationale:** Select UTRs with low-to-moderate predicted MRL and intermediate folding energies. This provides improvement headroom while keeping the structural constraint informative.<br>**Number and distribution:** Thirty 50-nt UTRs. Predicted MRL spans the 24th–27th percentiles and MFE the 21st–60th percentiles of the candidate pool.<br>**Sequence diversity:** Median pairwise Hamming distance of 37 out of 50 positions across 435 pairs. |

Table 9: **RNA folding energy**

| | |
|---|---|
| **Task description** | **Motivation:** RNA sequence determines secondary-structure stability. Lowering folding free energy while preserving GC content asks for a more favorable arrangement of nucleotides rather than simply increasing the fraction of G and C.<br>**Target:** Minimize predicted minimum folding free energy (MFE, kcal/mol).<br>**Constraint:** Preserve exact GC fraction within ±0.05 of the start. For 40-nt sequences, this permits a change of at most two GC bases. |
| **Trained predictors** | **Dataset:** 45,000 synthetic 40-nt RNAs labelled with ViennaRNA RNAfold [15]. These labels are computational thermodynamic predictions, not experimental measurements.<br>**Training split:** Random 90/10 training/validation split (seed 0); no separate local test partition.<br>**Model:** One-dimensional CNN approximating RNAfold MFE; exact nucleotide-count calculation for GC fraction.<br>**Performance:** Validation Spearman $\rho = 0.730$ for MFE; GC fraction is exact. |
| **Starting sequences** | **Selection rationale:** The first five are existing benchmark RNAs covering different folding energies and GC fractions; they predate the percentile-based selector, and the 25 added later were drawn with it. The target range is therefore wider here than on the other tasks.<br>**Number and distribution:** Thirty 40-nt RNAs. Target ranks span the 5th–82nd percentiles when lower MFE is counted as better; GC fractions span the 13th–68th percentiles of the candidate pool.<br>**Sequence diversity:** Median pairwise Hamming distance of 30 out of 40 positions across 435 pairs. |

#### 1.3 DNA Tasks

Table 10: **K562 enhancer activity**

| | |
|---|---|
| **Task description** | **Motivation:** Enhancers regulate where and how strongly genes are expressed. Cell-type-selective design requires increasing activity in the desired cellular context without broadly activating other cell types.<br>**Target:** Maximize K562 reporter activity ($\log_2$ RNA/DNA fold change).<br>**Constraints:** Preserve HepG2 and SK-N-SH activity, each within ±0.1 $\log_2$ RNA/DNA units of its starting value. |
| **Trained predictors** | **Dataset:** 798,064 lentiMPRA enhancer sequences measured in K562, HepG2, and SK-N-SH cells [9].<br>**Training split:** Random sequence split after deduplication: 638,452 training, 79,806 validation, and 79,806 test sequences per cell type.<br>**Model:** Three separate one-dimensional CNN regressors, one for each cell type.<br>**Performance:** Test Pearson $r = 0.862$ for K562, $r = 0.864$ for HepG2, and $r = 0.862$ for SK-N-SH. |
| **Starting sequences** | **Selection rationale:** Select low K562 activity to provide optimization headroom, with both constrained cell types between the 20th and 60th percentiles of their measured activity distributions. Favor sequence diversity.<br>**Number and distribution:** Thirty enhancers of 197–200 bp. K562 activity occupies the 0.1st–2.6th percentiles; HepG2 and SK-N-SH span the 42nd–50th and 42nd–51st percentiles, respectively. Percentiles use measured library values.<br>**Sequence diversity:** Median pairwise Hamming distance of 151 out of 200 bases, over the 406 of 435 pairs whose two starts have equal length. |

Table 11: **HepG2 enhancer activity**

| | |
|---|---|
| **Task description** | **Motivation:** This task seeks a HepG2-specific increase in enhancer activity while preventing a general increase across cellular contexts. Preserving activity in K562 and SK-N-SH distinguishes selective regulation from globally stronger enhancer activity.<br>**Target:** Maximize HepG2 reporter activity ($\log_2$ RNA/DNA fold change).<br>**Constraints:** Preserve K562 and SK-N-SH activity, each within ±0.1 $\log_2$ RNA/DNA units of its starting value. |
| **Trained predictors** | **Dataset:** The same 798,064-sequence three-cell-type lentiMPRA library [9].<br>**Training split:** Random 80/10/10 split after deduplication: 638,452 training, 79,806 validation, and 79,806 test sequences per predictor.<br>**Model:** Three independently trained one-dimensional CNN regressors, frozen during optimization.<br>**Performance:** Test Pearson $r = 0.864$ for HepG2, $r = 0.862$ for K562, and $r = 0.862$ for SK-N-SH. |
| **Starting sequences** | **Selection rationale:** Choose low HepG2 activity for improvement headroom while requiring the constrained cell types to lie in intermediate assay percentile ranges. Favor distinct sequences.<br>**Number and distribution:** Thirty enhancers of 199–200 bp. HepG2 target values occupy the 0.0th–3.1st percentiles; K562 and SK-N-SH span the 38th–49th and 40th–51st percentiles, respectively. Percentiles use measured library values.<br>**Sequence diversity:** Median pairwise Hamming distance of 149 out of 200 bases, over the 379 of 435 pairs whose two starts have equal length. |

Table 12: **SK-N-SH enhancer activity**

| | |
|---|---|
| **Task description** | **Motivation:** The enhancer should become more active in SK-N-SH neural-derived cells without increasing activity in K562 or HepG2. The task measures selective regulatory improvement rather than general enhancer strengthening.<br>**Target:** Maximize SK-N-SH reporter activity ($\log_2$ RNA/DNA fold change).<br>**Constraints:** Preserve K562 and HepG2 activity, each within ±0.1 $\log_2$ RNA/DNA units of its starting value. |
| **Trained predictors** | **Dataset:** 798,064 measured enhancer sequences with SK-N-SH, K562, and HepG2 readouts [9].<br>**Training split:** Random 80/10/10 sequence split: 638,452 training, 79,806 validation, and 79,806 test sequences per predictor.<br>**Model:** Three separate one-dimensional CNN regressors.<br>**Performance:** Test Pearson $r = 0.862$ for SK-N-SH, $r = 0.862$ for K562, and $r = 0.864$ for HepG2. |
| **Starting sequences** | **Selection rationale:** Choose low SK-N-SH activity while keeping K562 and HepG2 activity within intermediate percentile bands. Favor sequence diversity.<br>**Number and distribution:** Thirty enhancer sequences of 194–200 bp. SK-N-SH activities lie at the 0.0th–2.7th percentiles; K562 and HepG2 span the 42nd–51st and 41st–49th percentiles, respectively.<br>**Sequence diversity:** Median pairwise Hamming distance of 150 out of 200 bases, over the 325 of 435 pairs whose two starts have equal length. The rest differ in length and are not compared. |

Table 13: **Heat-shock-responsive promoter**

| | |
|---|---|
| **Task description** | **Motivation:** A useful heat-shock promoter should increase activity following heat exposure while maintaining approximately the same basal activity under control conditions. A constitutively stronger promoter does not satisfy this objective.<br>**Target:** Maximize predicted heat-shock inducibility, defined as heat-shock activity minus control activity, in PARM activity-score units.<br>**Constraints:** Preserve predicted control activity within ±0.2 score units of the start; at most 10 nucleotide substitutions. |
| **Trained predictors** | **Dataset:** K562 promoter-fragment MPRA under control conditions and 43°C heat shock, GSE301246, using the PARM architecture [4].<br>**Training split:** Five-fold promoter-fragment training with approximately 3.09 million training and 0.77 million validation fragments per fold. Starting promoters are selected from the official held-out test set.<br>**Model:** Separate PARM CNN ensembles for control and heat-shock activity. Inducibility is computed as their difference; substitution count is exact.<br>**Performance:** Held-out Pearson $r = 0.838$ for control, $r = 0.828$ for heat shock, and $r = 0.212$ for their predicted difference. |
| **Starting sequences** | **Selection rationale:** Require measurable basal activity between the 60th and 95th percentiles of eligible test promoters and low predicted inducibility below its 70th percentile. Also require stable predictions across folds, one promoter per gene, and separated loci.<br>**Number and distribution:** Eighty held-out promoters pass selection; the benchmark uses the first 30. Lengths range from 209 to 384 bp. Predicted control scores range from 1.043 to 1.622 and predicted inducibility from −0.485 to −0.110.<br>**Sequence diversity:** Pairwise Hamming distance is not defined without alignment because promoter lengths differ. |

Table 14: **Tissue-selective splice-site usage**

| | |
|---|---|
| **Task description** | **Motivation:** Splice-site usage can differ across tissues. The goal is to increase usage of an existing human splice site in liver while maintaining its usage in heart, brain, and testis. The canonical splice junction is protected from editing.<br>**Target:** Maximize predicted liver splice-site usage (fraction, 0–1).<br>**Constraints:** Preserve predicted heart, brain, and testis usage, each within ±0.05 of the start; at most 30 substitutions within a ±500-bp window. The splice-site core and its immediate flanking positions are fixed, leaving 993 mutable positions. |
| **Trained predictors** | **Dataset:** Released human splice-site models and tissue-specific usage data from Zeng and Li [22]. No predictor is retrained.<br>**Training split:** The selected sites are on Pangolin's held-out human chromosomes 1, 3, 5, 7, and 9, with one site per gene.<br>**Model:** Frozen Pangolin sequence model using 10,001 bp of genomic context. Predictions from five model replicates are averaged for each tissue. Edit count and protected positions are exact constraints.<br>**Performance:** Published Pangolin usage evaluation applies. The benchmark measures changes in predicted usage rather than reporting a newly trained oracle's test statistic. |
| **Starting sequences** | **Selection rationale:** Select real canonical splice sites with measured liver usage between 0.2 and 0.8 and predicted liver usage between 0.15 and 0.85, avoiding sites near zero or saturation. Require measurements in all four tissues and agreement among model replicates.<br>**Number and distribution:** Forty-eight distinct genes from held-out chromosomes. Observed starting liver usage ranges from 0.292 to 0.800 (median 0.658); predicted usage ranges from 0.168 to 0.850 (median 0.768).<br>**Sequence diversity:** Median pairwise Hamming distance of 7,467 out of 10,001 positions across 1,128 site pairs. Since the sites come from distinct genomic loci, this mainly describes differences among source sequences. |

### 2 Baseline

#### 2.1 Baseline Methods

We compare GRACE against baselines spanning five complementary optimization paradigms: direct gradient-based sequence optimization, penalty-based constrained optimization, evolutionary and stochastic search, black-box Bayesian optimization, and generative sequence design. All methods use the same frozen task oracles and starting sequences, and returned designs are evaluated using the same feasibility criteria. Table 15 summarizes the methods and their benchmark implementation; evaluation-budget matching is described separately in Section 6.

Table 15: **Baseline methods used for constrained biological sequence design.** The final column gives the central optimization principle and its implementation in our benchmark.

| Paradigm | Method | Optimization principle and implementation |
|---|---|---|
| Gradient-based | **Fast SeqProp** [14] | Directly maximizes the differentiable target oracle, $$\max_{\Theta} \; \bar{f}_T(\Theta),$$ using a continuous sequence parameterization and straight-through gradient estimation. Constraints are not used to determine the update direction; generated discrete sequences are subsequently evaluated under the benchmark feasibility criteria. |
| Evolutionary | **AdaLead** [20] | Adaptive population-based greedy search for biological sequence optimization. High-fitness sequences are retained as "leaders" and locally mutated to generate the next candidate population. In our benchmark, candidates are scored by the target oracle and the best sequence satisfying all task constraints is retained. |
| | **Directed evolution** [2] | Implements iterative mutation and selection in sequence space. From the current sequence $x_t$, neighboring mutants are evaluated and the highest-scoring feasible candidate is selected, $$x_{t+1} = \arg \max_{x \in \mathcal{N}(x_t),\, x \in \mathcal{F}} f_T(x),$$ where $\mathcal{F}$ denotes the feasible set defined by the task constraints. |
| Stochastic search | **Simulated annealing** [11] | Explores sequence space through stochastic mutations. Improving moves are accepted directly, while a move with target-score change $\Delta < 0$ is accepted with probability $$p_{\text{accept}} = \exp(\Delta/T),$$ where the temperature $T$ decreases during search. The best feasible sequence encountered is returned. |
| | **Random search** [5] | Samples mutated variants of the reference sequence without using gradients or an adaptive search model. Each candidate is evaluated by the same task oracles, and the highest-scoring feasible sequence found within the matched evaluation budget is returned. |
| Black-box Bayesian optimization | **Constrained Bayesian optimization** [3, 8] | Treats the target and constraint oracles as black-box functions. For each constraint, $$g_i(x) = \lvert f_{c_i}(x) - f_{c_i}(x_0) \rvert - \epsilon_i \le 0.$$ We fit Gaussian-process surrogates using BoTorch (`SingleTaskGP`) and use constrained `qLogExpectedImprovement`. Conceptually, acquisition favors both improvement and feasibility, $$\alpha(x) \propto \mathrm{EI}(x)\, P(g_1(x) \le 0, \ldots, g_m(x) \le 0)\,.$$ Because sequence space is discrete, acquisition is evaluated over pools of mutated candidate sequences represented by flattened one-hot encodings. |

Continued on next page

Table 15 – continued

| Paradigm | Method | Optimization principle and implementation |
|---|---|---|
| Generative | **Guided discrete diffusion** [10] | Uses a pretrained discrete sequence diffusion model as a generative prior and steers the reverse process using property gradients. We define the guidance score $$R(x) = f_T(x) - \lambda \sum_i \Big[ \lvert f_{c_i}(x) - f_{c_i}(x_0) \rvert - \epsilon_i \Big]_+ ,$$ and use its gradient to bias denoising toward higher target values while discouraging constraint violations. This follows the oracle-guided sequence diffusion paradigm introduced by NOS. |
| | **Prior-preserving guided diffusion** [10] | Adds an explicit likelihood-preservation term to property-guided diffusion. The guided transition is regularized toward the unguided sequence prior using $$\mathcal{L}_{\text{prior}} = \beta \, D_{\text{KL}}(q_{\text{guided}} \,\Vert\, q_{\text{prior}}) .$$ The parameter $\beta$ controls the trade-off between property optimization and remaining in high-likelihood regions of the pretrained sequence distribution. |
| | **Ctrl-DNA (For DNA regulatory tasks)** [6] | Constrained reinforcement learning for cell-type-specific regulatory DNA generation. Ctrl-DNA fine-tunes an *autoregressive genomic language model* to increase activity in a target cell type while controlling off-target activity. For the enhancer experiments, we apply the reported constrained-RL procedure using the HyenaDNA backbone and evaluate generated sequences under the same reference-centered feasibility criteria used for all methods. |

### 3 Constraint Behavior Along the Optimization Trajectory

The main results compare the best feasible designs returned by each method. Here, we examine the sequences explored during optimization. These are diagnostics rather than a second ranking: they ask whether GRACE's result follows from how it searches, or could have arisen by chance. A method is not required to lead on every one of them, and leading on one does not by itself establish an advantage, since each is a ratio computed within a method's own trajectory and is therefore sensitive to how many candidates that method proposes and how far it moves.

#### 3.1 Definitions

Let $\mathcal{H}$ denote the set of discrete sequences evaluated during a run, including the starting sequence $x_0$. Let $f$ denote the target score, oriented so that higher values are always better. For each constraint $j$, let $c_j$ denote the corresponding property, $c_{0,j} = c_j(x_0)$ its starting value, and $\epsilon_j$ its allowed tolerance. A sequence is feasible if $|c_j(x) - c_{0,j}| \le \epsilon_j$ for every constraint, together with any applicable mutation limit. We use $\tau = 10^{-6}$ to exclude negligible numerical changes when identifying target improvements.

**Feasibility.** Let $\mathcal{H}$ now denote every sequence a run generated. Feasibility is the percentage of those that both improve the target and satisfy all constraints:

$$\text{Feasibility} = 100 \times \frac{\#\{x \in \mathcal{H} : f(x) > f(x_0) + \tau \text{ and } x \text{ is feasible}\}}{\#\mathcal{H}}.$$

The denominator is the method's entire output for that run, not only its improving candidates. For one enhancer run it is 500 steps $\times K = 8$ samples per step, or 4,000 sequences; the table reports the median over the 60 runs of each task.

**Retention.** We measure how much of the best discovered target improvement remains available after enforcing the constraints:

$$\text{Retention} = 100 \times \frac{\max\{f(x) : x \in \mathcal{H},\ x \text{ is feasible}\} - f(x_0)}{\max\{f(x) : x \in \mathcal{H}\} - f(x_0)}.$$

The numerator is the best feasible improvement found in the run, and the denominator the best improvement found when the constraints are ignored, so retention asks how much of what a method discovered it is actually allowed to keep. Both extrema are taken over the same set $\mathcal{H}$ used for feasibility, so the measure is internal to a run and carries no information about the size of the gain. Runs that never improved the target have an undefined ratio and are dropped; the table reports the median over the remaining runs of each task.

**Violation severity.** For infeasible candidates, we measure how far the constraint values exceed their allowed tolerances:

$$V(x) = \frac{1}{m} \sum_{j=1}^{m} \max\left(0, \frac{|c_j(x) - c_{0,j}|}{\epsilon_j} - 1\right).$$

A feasible candidate has $V(x) = 0$. For a single constraint, $V(x) = 1$ means that its deviation is twice the allowed tolerance. We report the mean severity among candidates that both improve the target and violate at least one constraint; feasible candidates contribute $V = 0$ by definition and are excluded. Unlike feasibility and retention this is not a per-run median: the qualifying candidates from every run of a task are pooled into one mean, because a run that stays inside the band contributes none and a per-run average would weight a handful of candidates as heavily as thousands. For RNA folding, the GC-content constraint is recomputed from integer nucleotide counts to avoid numerical errors at the feasibility boundary.

**Sequence identity.** Two separate measures, both on the single design a run returns, never on its trajectory. *Identity to the start* is the percentage of positions of a returned design that match its own starting sequence, computed per run and reported as the median over the runs of a task; a run that found no feasible improvement returns $x_0$ and so scores 100%. *Pairwise identity* is the same percentage computed between two designs that share a starting sequence but differ in random seed. Every such pair within a start is formed, the pairs from all starts of a task are pooled, and the table reports their median. Designs from different starting sequences are never compared, and a pair is skipped if the two designs differ in length, which is why the variable-length tasks have no column for it.

Feasibility, retention and identity to the start are computed per run and summarized by their median. Violation severity is a mean over qualifying candidates pooled across runs, and pairwise identity a median over seed pairs pooled across starts.

#### 3.2 Feasibility

Among the search baselines, GRACE turns the largest share of what it generates into feasible improvements (Table 16): a median of 44.6%, against 16.5% for random search, 15.0% for directed evolution, 13.7% for AdaLead, 8.5% for simulated annealing and 1.3% for FastSeqProp. Simulated annealing is higher on the three enhancer tasks, where it also produces smaller feasible improvements.

#### 3.3 Retention

GRACE retains a median 95.8% of the improvement it discovers (Table 17), against 58.5% for AdaLead, 84.9% for directed evolution, 75.4% for simulated annealing, 73.4% for random search and 29.7% for FastSeqProp.

Table 16: **Trajectory feasibility (%), median over runs.** For each run, the percentage of all sequences it generated that both improved the target and satisfied every constraint at its benchmark tolerance, mutation budget included; the table reports the median of that percentage over runs. Dashes indicate unavailable trajectories. Guided diffusion leads this column for two reasons that have little to do with optimization. Starting sequences were deliberately chosen near the 25th percentile of the target, and diffusion draws from a prior trained on the whole library, so a typical sample already scores above the start: 96% of its sequences count as improvements without the method having to search. It also changes only 12–15% of positions, which keeps 72% of its samples inside the tolerance band. It proposes many small, mildly better sequences. What this column does not show is how much better: on amyloid nucleation diffusion gains 2.18 against GRACE's 4.56, and quadrupling its budget leaves that figure unchanged.

| Task | GRACE | AdaLead | Dir. evol. | Sim. ann. | Random | FastSeqProp | Cons. BO | Diffusion |
|---|---|---|---|---|---|---|---|---|
| Toehold switch | 23.9 | 1.4 | 2.7 | 11.3 | 15.0 | 1.4 | 26.2 | 55.6 |
| 5'UTR translation (MRL) | 90.5 | 14.5 | 14.2 | 39.0 | 39.0 | 0.8 | 54.8 | 87.8 |
| RNA folding (MFE) | 28.1 | 2.6 | 5.3 | 2.4 | 21.2 | 1.3 | 35.5 | 23.8 |
| K562 enhancer | 9.5 | 9.0 | 15.8 | 25.9 | 4.2 | 0.2 | 35.8 | 25.5 |
| HepG2 enhancer | 8.3 | 11.1 | 12.8 | 25.0 | 3.6 | 1.0 | 44.8 | 6.1 |
| SK-N-SH enhancer | 8.3 | 12.9 | 8.0 | 26.7 | 6.4 | 0.2 | 38.6 | 34.3 |
| Splice-site usage | 63.5 | 34.6 | 3.3 | 2.1 | 38.6 | 0.3 | – | – |
| Heat-shock promoter | 68.4 | 4.4 | 43.2 | 0.0 | 2.0 | 1.0 | 45.5 | – |
| TEM-1 activity | 83.9 | 87.1 | 39.6 | 0.6 | – | 5.2 | – | 90.4 |
| TEM-1 solubility | 83.5 | 27.7 | 51.0 | 0.9 | – | 5.9 | – | 94.0 |
| Catalytic turnover | 26.0 | 6.5 | 6.4 | 5.8 | 9.9 | 1.2 | 41.9 | 53.9 |
| Protein solubility | 28.4 | 26.1 | 25.6 | 4.9 | 18.0 | 2.5 | 65.8 | 83.3 |
| Amyloid nucleation | 60.7 | 30.4 | 24.9 | 38.9 | 40.5 | 6.2 | 80.0 | 93.5 |
| Mini-protein stability | 78.5 | 34.2 | 22.8 | 23.0 | 30.0 | 4.2 | 72.1 | 95.0 |
| *Median* | 44.6 | 13.7 | 15.0 | 8.5 | 16.5 | 1.3 | 44.8 | 69.5 |

Table 17: **Constraint retention (%), median over runs.** Percentage of discovered target improvement retained after enforcing all constraints. Runs that never improved the target are dropped, since the ratio is undefined for them. Dashes indicate unavailable trajectories; column medians cover only the tasks a method was run on. Retention is a ratio within each method's own trajectory and says nothing about the size of the gain: guided diffusion retains 100% of a 0.03 improvement on TEM-1 activity. Absolute improvements are reported in the main text.

| Task | GRACE | AdaLead | Dir. evol. | Sim. ann. | Random | FastSeqProp | Cons. BO | Diffusion |
|---|---|---|---|---|---|---|---|---|
| Toehold switch | 88.1 | 48.0 | 35.2 | 60.8 | 66.1 | 11.4 | 73.9 | 97.6 |
| 5'UTR translation (MRL) | 100.0 | 92.5 | 88.4 | 89.7 | 95.8 | 73.1 | 100.0 | 100.0 |
| RNA folding (MFE) | 55.3 | 17.0 | 13.7 | 7.3 | 52.7 | 8.0 | 58.7 | 57.2 |
| K562 enhancer | 95.2 | 45.9 | 93.9 | 86.3 | 25.7 | 8.4 | 91.5 | 93.7 |
| HepG2 enhancer | 91.3 | 38.5 | 81.3 | 89.1 | 19.8 | 3.8 | 94.5 | 33.5 |
| SK-N-SH enhancer | 93.1 | 37.1 | 76.6 | 93.1 | 30.4 | 3.8 | 92.1 | 82.7 |
| Splice-site usage | 86.2 | 69.0 | 16.9 | 33.2 | 93.0 | 23.5 | – | – |
| Heat-shock promoter | 96.4 | 5.7 | 88.5 | 0.6 | 5.6 | 0.3 | 48.9 | – |
| TEM-1 activity | 100.0 | 100.0 | 100.0 | 14.5 | – | 35.9 | – | 100.0 |
| TEM-1 solubility | 99.7 | 42.4 | 100.0 | 7.8 | – | 38.0 | – | 100.0 |
| Catalytic turnover | 99.0 | 83.5 | 90.2 | 74.5 | 80.8 | 90.1 | 85.5 | 97.8 |
| Protein solubility | 99.4 | 94.2 | 99.6 | 77.7 | 89.8 | 90.4 | 100.0 | 100.0 |
| Amyloid nucleation | 95.2 | 94.7 | 77.7 | 76.3 | 100.0 | 74.8 | 100.0 | 100.0 |
| Mini-protein stability | 97.4 | 96.4 | 55.8 | 93.7 | 98.5 | 79.9 | 100.0 | 100.0 |
| *Median* | 95.8 | 58.5 | 84.9 | 75.4 | 73.4 | 29.7 | 92.1 | 98.9 |

The difference is clearest for FastSeqProp, which optimizes the target without explicitly handling the constraints. It often discovers high-scoring sequences, but most of that improvement is lost once infeasible candidates are excluded. GRACE instead keeps nearly all of what it finds.

Table 18: **Constraint violation when infeasible.** Mean normalised excess $V(x) = \text{mean}_j \max(0, |c_j - c_{0,j}|/\epsilon_j - 1)$, pooled across all runs of a task over the logged candidates that improve the target and fall outside the band; feasible candidates have $V = 0$ and are excluded. $V = 1$ means twice the tolerance. Lower is better. Constrained BO and guided diffusion are comparable to GRACE on this measure, consistent with the small edits they propose.

| Task | GRACE | AdaLead | Dir. evol. | Sim. ann. | Random | FastSeqProp | Cons. BO | Diffusion |
|---|---|---|---|---|---|---|---|---|
| Toehold switch | 1.7 | 22.1 | 21.5 | 6.4 | 4.2 | 34.2 | 2.8 | 0.9 |
| 5'UTR translation (MRL) | 0.2 | 1.3 | 1.0 | 1.0 | 0.8 | 2.9 | 0.4 | 0.1 |
| RNA folding (MFE) | 1.0 | 5.7 | 5.6 | 5.7 | 0.8 | 8.8 | 0.3 | 0.4 |
| K562 enhancer | 1.9 | 3.9 | 1.6 | 1.3 | 5.4 | 69.6 | 1.1 | 1.1 |
| HepG2 enhancer | 2.2 | 3.0 | 1.7 | 1.0 | 5.9 | 95.8 | 1.0 | 2.2 |
| SK-N-SH enhancer | 2.2 | 2.9 | 2.4 | 1.2 | 5.7 | 87.4 | 1.0 | 0.9 |
| Splice-site usage | 0.3 | 0.3 | 0.7 | 0.6 | 0.2 | 2.2 | – | – |
| Heat-shock promoter | 0.3 | 2.1 | 0.9 | 10.1 | 5.8 | 8.3 | 1.4 | – |
| TEM-1 activity | 0.1 | 0.1 | – | 1.6 | – | 0.3 | – | – |
| TEM-1 solubility | 0.1 | 0.7 | – | 1.8 | – | 1.9 | – | 0.1 |
| Catalytic turnover | 2.3 | 8.7 | 7.8 | 9.7 | 6.2 | 13.1 | 2.3 | 0.7 |
| Protein solubility | 3.5 | 4.1 | 3.8 | 6.1 | 4.6 | 9.8 | 2.3 | 1.1 |
| Amyloid nucleation | 1.1 | 2.1 | 3.4 | 1.7 | 1.5 | 4.2 | 0.9 | 0.9 |
| Mini-protein stability | 1.0 | 3.3 | 6.9 | 2.5 | 2.3 | 7.6 | 1.2 | 0.9 |
| *Median* | 1.1 | 3.0 | 2.9 | 1.7 | 4.4 | 8.6 | 1.1 | 0.9 |

#### 3.4 Violation Severity

Table 19: **Pairwise identity among returned designs (%).** Sequence identity between designs generated from the same starting sequence using different random seeds. Every within-start seed pair is formed and the pairs from all starts are pooled; the table reports their median. Lower identity indicates greater diversity among returned designs. A cell requires at least two seeds per start, so splice-site usage is excluded and methods run at a single seed on a task are left blank. Constrained BO is taken from the matched-budget campaign (10 starts × 2 seeds), the only BO runs with more than one seed per start; its iteration count is fixed, so both run sets spend the same number of target calls.

| Task | GRACE | AdaLead | Dir. evol. | Sim. ann. | Random | FastSeqProp | Cons. BO | Diffusion |
|---|---|---|---|---|---|---|---|---|
| Toehold switch | 86.7 | 46.7 | 100.0 | 33.3 | 36.7 | 100.0 | 46.7 | 83.3 |
| 5'UTR translation (MRL) | 72.0 | 46.0 | 100.0 | 26.0 | 40.0 | 100.0 | 51.0 | 92.0 |
| RNA folding (MFE) | 85.0 | 67.5 | 100.0 | 35.0 | 47.5 | 97.5 | 53.8 | 92.5 |
| K562 enhancer | 72.0 | 89.0 | 100.0 | 24.5 | 40.0 | 100.0 | 62.0 | – |
| HepG2 enhancer | 76.5 | 91.0 | 100.0 | 24.5 | 37.0 | 99.5 | 58.2 | 75.5 |
| SK-N-SH enhancer | 71.5 | 89.2 | 100.0 | 25.5 | 31.5 | 99.5 | 62.8 | – |
| Splice-site usage | – | – | – | – | – | – | – | – |
| Heat-shock promoter | 98.2 | 94.2 | 100.0 | 97.3 | – | 100.0 | – | – |
| TEM-1 activity | 83.0 | 68.6 | 99.3 | 95.8 | 85.3 | 91.2 | 96.1 | 96.8 |
| TEM-1 solubility | 70.2 | 81.9 | 99.3 | 87.9 | 78.6 | 87.4 | 80.9 | 98.2 |
| Catalytic turnover | 55.0 | 22.5 | 100.0 | 5.0 | – | 82.5 | 56.2 | 60.0 |
| Protein solubility | 55.0 | 22.5 | 100.0 | 10.0 | – | 40.0 | 27.5 | 60.0 |
| Amyloid nucleation | 75.0 | 20.0 | 100.0 | 5.0 | – | 60.0 | 12.5 | 65.0 |
| Mini-protein stability | 83.7 | 11.6 | 100.0 | 7.0 | – | 81.4 | 24.4 | 58.1 |
| *Median* | 75.0 | 67.5 | 100.0 | 25.5 | 40.0 | 97.5 | 55.0 | 79.4 |

GRACE also tends to produce smaller constraint violations when its improving candidates are infeasible. It has the lowest average violation severity among the comparably-scaled search baselines on 9 of the 14 tasks (Table 18).

Its median severity is 1.05, compared with 2.99 for AdaLead, 2.88 for directed evolution, 1.75 for simulated annealing,

4.40 for random search and 8.58 for FastSeqProp. The toehold switch shows a particularly large difference: GRACE exceeds the constraint limits by an average of 1.73 tolerance units, compared with 22.08 for AdaLead and 34.24 for FastSeqProp.

Together with the feasibility and retention results, this suggests that GRACE generally limits constraint violations while continuing to improve the target property.

##### 3.5 Sequence Identity

GRACE does not preserve constraints simply by making small edits.

Across seeds from the same start (Table 19), GRACE has a median pairwise identity of 75.0%, against 67.5% for AdaLead, 100.0% for directed evolution, 55.0% for constrained BO and 79.4% for guided diffusion. The pattern is task-dependent: on the four CNN protein tasks GRACE converges to more similar designs than AdaLead (55.0–83.7% versus 11.6–22.5%), while on the three enhancer tasks it is the more diverse of the two (71.5–76.5% versus 89.0–91.0%). Multiple starting sequences are therefore the more reliable route to a broad set of distinct designs.

#### 4 Robustness to Constraint Tolerances

Table 20: **Best feasible target improvement across constraint tolerances.** Entries are median improvements over 15 runs (five starting sequences and three seeds). Constraint tolerances are scaled relative to their benchmark values; evaluation and mutation budgets are fixed. Higher is better. Bold indicates the best method for each task and tolerance, including the two GRACE ablations. **This sweep predates the 30-start campaign** and retains the earlier five-start, three-seed configuration, so its absolute values are not directly comparable with Table 21, which uses 30 starting sequences at two seeds. The two differ most on the K562 enhancer, where the smaller start set reverses the ordering of GRACE and ALM-only; the comparison to use is Table 21. What this table is for is the *within-configuration* trend across tolerances, which is unaffected.

| Domain | Task | Method | 0.5× | 0.75× | 1× | 1.5× | 2× |
|---|---|---|---|---|---|---|---|
| **Protein** | **Protein solubility** | **GRACE** | **281.32** | **282.33** | 279.85 | **283.12** | **280.35** |
| | | Projection only | 261.83 | 255.02 | 274.99 | 277.70 | 272.10 |
| | | ALM only | 275.67 | 279.62 | **285.66** | 280.16 | 277.43 |
| | | AdaLead | 172.91 | 172.91 | 175.37 | 175.37 | 177.79 |
| | | Directed evolution | 237.59 | 237.59 | 237.59 | 237.59 | 237.59 |
| | | Simulated annealing | 185.87 | 185.87 | 213.62 | 213.62 | 233.43 |
| | | FastSeqProp | 276.62 | 276.62 | 276.62 | 276.62 | 278.36 |
| **Protein** | **Amyloid nucleation** | **GRACE** | **4.60** | **4.57** | 4.60 | 4.67 | **4.74** |
| | | Projection only | 4.59 | 4.54 | **4.62** | **4.70** | **4.74** |
| | | ALM only | 4.50 | 4.53 | 4.58 | 4.65 | **4.74** |
| | | AdaLead | 3.37 | 3.37 | 3.71 | 3.71 | 3.83 |
| | | Directed evolution | 2.24 | 2.24 | 2.67 | 3.86 | 3.94 |
| | | Simulated annealing | 2.49 | 2.49 | 2.76 | 2.76 | 2.76 |
| | | FastSeqProp | 3.63 | 3.63 | 3.68 | 4.26 | 4.31 |
| **Protein** | **Mini-protein stability** | **GRACE** | **5.83** | **5.82** | 5.82 | **5.88** | 5.88 |
| | | Projection only | 5.81 | 5.80 | 5.80 | 5.87 | 5.87 |
| | | ALM only | 5.82 | 5.80 | **5.83** | 5.87 | **5.89** |
| | | AdaLead | 3.95 | 3.95 | 4.05 | 4.16 | 4.38 |
| | | Directed evolution | 3.02 | 3.02 | 3.20 | 3.20 | 3.37 |
| | | Simulated annealing | 3.66 | 3.66 | 3.93 | 4.11 | 4.13 |
| | | FastSeqProp | 3.65 | 3.65 | 4.28 | 4.28 | 4.30 |
| **RNA** | **Toehold switch** | **GRACE** | **0.98** | **0.98** | **0.99** | **1.01** | **1.02** |
| | | Projection only | 0.91 | 0.92 | 0.95 | 0.85 | 0.91 |
| | | ALM only | 0.93 | 0.93 | 0.93 | 0.97 | 0.94 |
| | | AdaLead | 0.57 | 0.62 | 0.62 | 0.67 | 0.70 |
| | | Directed evolution | 0.17 | 0.45 | 0.45 | 0.45 | 0.57 |
| | | Simulated annealing | 0.62 | 0.63 | 0.63 | 0.65 | 0.71 |

| | | | | | | | |
|---|---|---|---|---|---|---|---|
| | | FastSeqProp | 0.14 | 0.14 | 0.14 | 0.27 | 0.29 |
| **RNA** | **5′UTR translation (MRL)** | **GRACE** | 2.60 | 2.64 | **2.72** | 2.68 | 2.69 |
| | | Projection only | 2.50 | 2.52 | 2.60 | 2.69 | 2.70 |
| | | ALM only | **2.63** | **2.65** | 2.66 | 2.71 | 2.81 |
| | | AdaLead | 2.35 | 2.37 | 2.51 | 2.70 | 2.77 |
| | | Directed evolution | 2.09 | 2.21 | 2.59 | **2.80** | **2.84** |
| | | Simulated annealing | 2.08 | 2.13 | 2.16 | 2.30 | 2.34 |
| | | FastSeqProp | 1.94 | 2.00 | 2.04 | 2.07 | 2.18 |
| **RNA** | **RNA folding (MFE)** | **GRACE** | **6.06** | **6.81** | **6.69** | **8.00** | **8.86** |
| | | Projection only | 3.92 | 2.24 | 2.24 | 4.74 | 4.87 |
| | | ALM only | 1.61 | 1.67 | 1.67 | 3.56 | 4.34 |
| | | AdaLead | 2.86 | 2.97 | 2.97 | 5.02 | 5.59 |
| | | Directed evolution | 2.67 | 2.71 | 2.71 | 5.61 | 5.69 |
| | | Simulated annealing | 2.02 | 2.04 | 2.04 | 3.55 | 4.77 |
| | | FastSeqProp | 1.61 | 1.67 | 1.67 | 3.56 | 4.34 |
| **DNA** | **K562 enhancer** | **GRACE** | 8.21 | 8.21 | 8.75 | 8.62 | 8.52 |
| | | Projection only | 4.46 | 6.65 | 6.38 | 6.86 | 8.07 |
| | | ALM only | **9.55** | **9.69** | **9.68** | **9.87** | **10.00** |
| | | AdaLead | 1.10 | 1.43 | 1.72 | 1.74 | 1.91 |
| | | Directed evolution | 6.22 | 6.80 | 7.27 | 7.84 | 6.73 |
| | | Simulated annealing | 3.41 | 4.44 | 4.57 | 5.14 | 5.58 |
| | | FastSeqProp | 0.66 | 1.14 | 1.36 | 1.51 | 1.65 |
| **DNA** | **HepG2 enhancer** | **GRACE** | **7.87** | **8.13** | **7.86** | **8.25** | **8.18** |
| | | Projection only | 2.20 | 2.66 | 3.64 | 3.78 | 2.98 |
| | | ALM only | 7.76 | 7.56 | 7.52 | 7.96 | 7.92 |
| | | AdaLead | 0.53 | 0.77 | 0.77 | 0.82 | 1.01 |
| | | Directed evolution | 4.32 | 4.20 | 5.12 | 4.16 | 4.18 |
| | | Simulated annealing | 3.15 | 3.06 | 3.49 | 3.71 | 4.33 |
| | | FastSeqProp | 0.41 | 0.76 | 0.89 | 1.04 | 1.29 |
| **DNA** | **SK-N-SH enhancer** | **GRACE** | **10.25** | **10.43** | **10.50** | **10.81** | **11.15** |
| | | Projection only | 7.85 | 5.70 | 4.11 | 4.21 | 3.36 |
| | | ALM only | 9.61 | 9.66 | 9.58 | 9.35 | 10.23 |
| | | AdaLead | 0.79 | 0.88 | 1.01 | 1.12 | 1.26 |
| | | Directed evolution | 5.67 | 6.13 | 7.06 | 6.47 | 6.94 |
| | | Simulated annealing | 4.46 | 4.65 | 5.29 | 5.96 | 5.83 |
| | | FastSeqProp | 0.75 | 0.75 | 1.07 | 1.14 | 1.70 |

### 5 Component Ablation

GRACE has two parts: a null-space projection that removes the component of the target gradient which would push an active constraint out of its band, and an augmented Lagrangian that corrects whatever violation remains. Table 21 runs each alone on every task.

Neither is sufficient, and they fail differently. Projection only collapses where the constraint geometry is hard: no feasible improvement in any of the 15 RNA-folding runs, and 1–4% of GRACE's gain on the three enhancers. ALM only tracks GRACE on most tasks but returns nothing on RNA folding and gives up a fifth to a third of the enhancer improvement. The full method is the only arm that never collapses: it returns a feasible improvement in every run on 12 of the 13 tasks with paired gain data, and in 87% of runs on the other.

The ablations are not uniformly worse. Projection only gives the largest gain on catalytic turnover, and ALM only ties GRACE on both TEM-1 tasks and edges it on the 5′UTR. What separates the full method is not winning every cell but never collapsing in any of them, which is what the per-task success column shows.

Table 21: **Component ablation.** Projection only keeps the active-set null-space projection and drops the multipliers; ALM only keeps the augmented Lagrangian and drops the projection. *Best feasible improvement* is the median improvement over the start in that task's own oracle units, so it is comparable only within a task. *Success* is the percentage of runs returning any feasible improvement. Both use the 274 cells where all three arms ran the same start and seed. Trajectory feasibility and violation severity are defined in Section 3. Bold marks the best arm per task. The heat-shock gain and success cells are omitted because its ablation arms were not run under the same restart protocol (Section 8).

| Domain | Task | Arm | Best feasible improvement | Success (%) | Feasibility (%) | Violation $V$ |
|---|---|---|---|---|---|---|
| **RNA** | Toehold switch | GRACE | **0.95** | **100** | **23.9** | **1.73** |
| | | Projection only | 0.81 | 80 | 11.4 | 6.90 |
| | | ALM only | **0.95** | 93 | 18.4 | 2.82 |
| | 5'UTR translation (MRL) | GRACE | 2.63 | **100** | **90.5** | 0.24 |
| | | Projection only | 2.58 | 87 | 80.1 | 1.01 |
| | | ALM only | **2.65** | **100** | 73.9 | **0.18** |
| | RNA folding (MFE) | GRACE | **8.20** | **87** | **28.1** | **1.00** |
| | | Projection only | 0.00 | 0 | 1.5 | 2.76 |
| | | ALM only | 0.00 | 40 | 1.3 | 8.54 |
| **DNA** | K562 enhancer | GRACE | **11.76** | **100** | **9.5** | **1.91** |
| | | Projection only | 0.22 | 60 | 1.3 | 8.91 |
| | | ALM only | 9.18 | **100** | 4.7 | 3.95 |
| | HepG2 enhancer | GRACE | **9.71** | **100** | **8.3** | **2.22** |
| | | Projection only | 0.05 | 67 | 2.8 | 22.81 |
| | | ALM only | 6.58 | **100** | 3.7 | 9.09 |
| | SK-N-SH enhancer | GRACE | **12.34** | **100** | **8.3** | **2.21** |
| | | Projection only | 0.49 | 93 | 3.2 | 20.45 |
| | | ALM only | 8.49 | **100** | 5.4 | 9.41 |
| | Splice-site usage | GRACE | **0.09** | **100** | **63.5** | **0.27** |
| | | Projection only | 0.07 | 98 | 29.8 | 0.41 |
| | | ALM only | 0.07 | 98 | 59.7 | 0.32 |
| **Protein** | TEM-1 activity | GRACE | **0.28** | **100** | 83.9 | **0.06** |
| | | Projection only | 0.06 | **100** | 5.2 | 0.52 |
| | | ALM only | **0.28** | **100** | **91.7** | 0.09 |
| | TEM-1 solubility | GRACE | **0.27** | **100** | **83.5** | **0.15** |
| | | Projection only | 0.09 | **100** | 6.0 | 1.82 |
| | | ALM only | **0.27** | **100** | 76.2 | 0.16 |
| | Catalytic turnover | GRACE | 11.67 | **100** | 26.0 | **2.26** |
| | | Projection only | **13.10** | **100** | 6.8 | 6.19 |
| | | ALM only | 11.82 | **100** | **26.7** | **2.26** |
| | Protein solubility | GRACE | **278.19** | **100** | **28.4** | 3.46 |
| | | Projection only | 266.35 | **100** | 12.7 | 8.24 |
| | | ALM only | 274.71 | **100** | 28.3 | **3.44** |
| | Amyloid nucleation | GRACE | **4.56** | **100** | **60.7** | **1.10** |
| | | Projection only | **4.56** | **100** | 36.8 | 1.51 |
| | | ALM only | 4.52 | **100** | 55.8 | 1.14 |
| | Mini-protein stability | GRACE | **5.81** | **100** | **78.5** | 1.00 |
| | | Projection only | 5.35 | **100** | 14.4 | 2.27 |
| | | ALM only | 5.80 | **100** | 69.1 | **0.95** |

### 6 Fairness of Baseline Comparison

All methods use the same starting sequences, frozen oracles, constraints and final selection rule, and every run returns the best feasible discrete sequence encountered, or the starting sequence if none was found.

Budgets are matched on predictor calls. A GRACE run with $T$ steps, $K$ samples per step and $m$ constraint oracles spends

$$G = T\,K\,(1 + m) \tag{1}$$

calls, and every baseline is given the same $G$ on the same task. Each implementation takes its budget in its own units – candidates, draws, steps or an evaluation cap – so the value passed to each one is set to whatever makes its call count equal $G$, and the achieved count is read back from each run's own counter rather than assumed.

Table 22: Predictor calls per run, $G = TK(1 + m)$, applied to every method on that task.

| **Task** | $T$ | $K$ | $1 + m$ | $G$ |
|---|---|---|---|---|
| Toehold switch | 500 | 8 | 2 | 8,000 |
| 5′UTR translation (MRL) | 800 | 16 | 2 | 25,600 |
| RNA folding (MFE) | 500 | 16 | 2 | 16,000 |
| Enhancers (K562, HepG2, SK-N-SH) | 500 | 8 | 3 | 12,000 |
| TEM-1 activity / solubility | 500 | 8 | 4 | 16,000 |
| Catalytic turnover | 800 | 16 | 2 | 25,600 |
| Protein solubility | 800 | 16 | 2 | 25,600 |
| Amyloid nucleation | 400 | 8 | 2 | 6,400 |
| Mini-protein stability | 800 | 16 | 2 | 25,600 |
| Heat-shock promoter | 300 | 8 | 3 | 15,000 |
| Splice-site usage | 300 | 2 | – | 900 queries |

One Pangolin query returns all four tissues at once, so splicing is counted in sequence queries rather than separate oracle calls.

Constrained BO is the exception. Its implementation caps the number of acquisition iterations at $\min(\text{budget} - 20,\ 180)$, because refitting the exact GP dominates its cost, so it evaluates 150–200 candidates whatever budget it is given. We therefore run it at its published setting and report the candidates it actually evaluated alongside its wall-clock time, rather than describing it as budget-matched.

### 7 GRACE Implementation Details

#### 7.1 Parameters

Hyperparameters were selected on the first 5 of the 30 starting sequences; the remaining 25 are held out and never used for tuning. The criterion was whether a setting leaves GRACE stuck at the starting sequence. $\rho$ is the only coefficient that varies across tasks, and the only one re-selected in the final round (RNA folding, 10 $\rightarrow$ 100).

Table 23: GRACE hyperparameters. $K$ Gumbel-Softmax samples per step, $\eta$ the Adam learning rate on the logits, $\rho$ the augmented-Lagrangian penalty. Shared across all tasks: active-set threshold $\gamma = 0.3$, projection damping $\delta = 10^{-6}$, dual step $\eta_\lambda = 0.05$ (0.02 on TEM-1 and the enhancers), multiplier clipping $\lambda_i \in [0, 100]$.

| **Domain** | **Task** | $K$ | **Steps** | $\eta$ | $\rho$ |
|---|---|---|---|---|---|
| RNA | Toehold switch | 8 | 500 | 0.20 | 10 |
| RNA | 5′UTR translation (MRL) | 16 | 800 | 0.25 | 10 |
| RNA | RNA folding (MFE) | 16 | 500 | 0.20 | 100 |
| DNA | Enhancers (K562, HepG2, SK-N-SH) | 8 | 500 | 0.01 | 10 |
| DNA | Splice-site usage | 2 | 300 | 0.05 | 10 |
| DNA | Heat-shock promoter | 8 | 300 | 0.10 | 1.0 |
| Protein | TEM-1 activity / solubility | 8 | 500 | 0.03 | 10 |
| Protein | Catalytic turnover | 16 | 800 | 0.10 | 0.1 |
| Protein | Protein solubility | 16 | 800 | 0.20 | 1.0 |
| Protein | Amyloid nucleation | 8 | 400 | 0.10 | 0.5 |
| Protein | Mini-protein stability | 16 | 800 | 0.10 | 0.1 |

Logits are initialized from the one-hot encoding of the starting sequence, scaled by four. Each step draws $K$ candidates by Gumbel–Softmax with a straight-through estimator, Gumbel noise scaled by 0.5, temperature annealed as $\tau_t = \max(0.3,\ 0.3^{t/T})$; target and constraint values are averaged over the $K$ samples. Constraint references $c_i^0$ are computed once from the starting sequence. Feasibility and final selection are evaluated on decoded discrete sequences: each run returns the highest-scoring feasible sequence encountered, or the starting sequence if none was found. The same selection rule applies to every baseline.

#### 7.2 Dual Updates

The multipliers $\lambda_i$ in the augmented-Lagrangian term of Algorithm 1 are not fixed. Each starts at $\lambda_i = 0$ and is updated once per step, after the parameter update, by gradient ascent on the constraint residual:

$$\lambda_i \ \leftarrow\ \mathrm{clip}(\lambda_i + \eta_\lambda\, h_i(\Theta),\ 0,\ \lambda_{\max}), \qquad h_i(\Theta) = \left|\bar{f}_{c_i}(\Theta) - c_i^0\right| - \epsilon_i, \tag{2}$$

with dual step $\eta_\lambda$ and $\lambda_{\max} = 100$. Clipping at zero keeps the multipliers non-negative, as the inequality form requires, and $\lambda_{\max}$ bounds the linear term so that a persistently violated constraint cannot overwhelm the target gradient.

The update is driven by the signed residual $h_i$ rather than the clipped violation $v_i$, so a constraint sitting inside its band ($h_i < 0$) relaxes its own multiplier instead of holding whatever value it reached earlier. This is safe because $\lambda_i$ multiplies $v_i$, which is zero whenever constraint $i$ is satisfied: a multiplier that has grown large has no effect while the constraint is respected, and acts only once the band is crossed. $\eta_\lambda = 0.05$ on all tasks except TEM-1 and the three enhancers, where it is 0.02 (Table 23). The penalty coefficient $\rho$ is fixed per task and is not adapted during a run.

**Two implementation details.** The three enhancer tasks run through a separate DNA optimizer whose dual update uses the clipped violation in place of the signed residual, $\lambda_i \leftarrow \mathrm{clip}(\lambda_i + \eta_\lambda v_i,\ 0,\ \lambda_{\max})$, so there the multipliers are non-decreasing within a run. On the remaining eleven tasks the residual in Eq. (2) is evaluated against a single scalar tolerance rather than the per-constraint $\epsilon_i$ that defines $v_i$. The two agree wherever a task has a single learned constraint, and differ only for the Hamming mutation budget on TEM-1, heat shock and splicing, whose $\epsilon_i$ is larger; there that multiplier rises faster than Eq. (2) states, which changes nothing unless the budget is actually exceeded.

#### 7.3 GRACE Algorithm

#### 7.4 Notation

Let $x \in \mathcal{A}^L$ denote a sequence of length $L$ over an alphabet of size $A$ ($A = 4$ for nucleotides and 20 for amino acids). GRACE optimizes continuous logits $\theta \in \mathbb{R}^{A\times L}$, whose column-wise softmax defines a distribution over sequence symbols.

Each task specifies a differentiable target predictor $f$ (larger is better) and $m$ differentiable constraint predictors $c_1, \ldots, c_m$. Given a starting sequence $x_0$, a design is feasible if

$$|c_j(x) - c_j(x_0)| \leq \epsilon_j, \qquad j = 1, \ldots, m, \tag{3}$$

where $\epsilon_j$ is the constraint tolerance. Mutation budgets, when applicable, are included as Hamming-distance constraints.

#### 7.5 Optimization Procedure

GRACE combines three components for constrained sequence optimization.

**Straight-through relaxation.** Gumbel–Softmax sampling enables gradient-based optimization of discrete sequences, using hard one-hot representations in the forward pass and soft gradients in the backward pass.

**Active-set null-space projection.** The target gradient is projected onto the approximate null space of constraint gradients whose deviations exceed a fraction $\kappa$ of their tolerances, preserving progress along feasible directions.

**Augmented-Lagrangian correction.** Linear dual and quadratic penalties correct residual constraint violations, with dual variables updated throughout optimization.

The complete procedure is presented in Algorithm 1.

**Algorithm 1** GRACE: Constrained Biological Sequence Optimization

**Require:** Starting sequence $x_0$; target predictor $f$; constraint predictors $c_1, \dots, c_m$; tolerances $\epsilon_1, \dots, \epsilon_m$; iterations $T$; samples per iteration $K$; learning rate $\eta$; penalty $\rho$; dual learning rate $\eta_\lambda$; active-set threshold $\kappa = 0.3$; ridge $\delta = 10^{-6}$

**Ensure:** Best feasible discrete sequence

$\theta \leftarrow 4\,\mathrm{onehot}(x_0)$
$c_{0,j} \leftarrow c_j(x_0), \quad \lambda_j \leftarrow 0$ for all $j$, $\quad \mathcal{S} \leftarrow \{x_0\}$
**for** $t = 0, \dots, T-1$ **do**
  $\tau_t \leftarrow \max(0.3,\ 0.3^{t/T})$
  **for** $k = 1, \dots, K$ **do**
    $G^{(k)} \sim \mathrm{Gumbel}(0,1)^{A \times L}$
    $P^{(k)} \leftarrow \mathrm{softmax}\big((\theta + 0.5G^{(k)})/\tau_t\big)$
    $X^{(k)} \leftarrow \mathrm{onehot}(\arg\max P^{(k)}) + P^{(k)} - \mathrm{sg}[P^{(k)}]$
  **end for**
  $\bar{f} \leftarrow K^{-1} \sum_k f(X^{(k)}), \quad \bar{c}_j \leftarrow K^{-1} \sum_k c_j(X^{(k)})$
  $g \leftarrow \nabla_\theta \bar{f}$
  {1. Active-set null-space projection}
  $\mathcal{A}_t \leftarrow \{j : |\bar{c}_j - c_{0,j}| \geq \kappa\epsilon_j\}$
  **if** $\mathcal{A}_t \neq \emptyset$ **then**
    $J \leftarrow [\nabla_\theta \bar{c}_j]_{j \in \mathcal{A}_t}$ {constraint gradients as rows}
    $d \leftarrow g - J^\top (JJ^\top + \delta I)^{-1} Jg$
  **else**
    $d \leftarrow g$
  **end if**
  {2. Augmented-Lagrangian correction}
  $v_j \leftarrow \max(0, |\bar{c}_j - c_{0,j}| - \epsilon_j)$
  $\Phi \leftarrow \sum_{j=1}^{m} \left(\lambda_j v_j + \frac{\rho}{2} v_j^2\right)$
  {3. Parameter and dual updates}
  $\theta \leftarrow \theta + \eta\,\mathrm{Adam}\big(d - \nabla_\theta \Phi\big)$
  $\lambda_j \leftarrow \mathrm{clip}\big(\lambda_j + \eta_\lambda(|\bar{c}_j - c_{0,j}| - \epsilon_j),\ 0,\ 100\big)$
  **if** $t \bmod 3 = 0$ **then**
    $\mathcal{S} \leftarrow \mathcal{S} \cup \{\mathrm{decode}(\theta)\}$
  **end if**
**end for**
$\mathcal{S} \leftarrow \mathcal{S} \cup \{\mathrm{decode}(\theta)\}$
**return** $\arg \max_{\substack{x \in \mathcal{S} \\ x \text{ feasible}}} f(x)$

Here, sg denotes the stop-gradient operator, and decode$(\theta)$ applies a position-wise argmax to obtain a discrete sequence. Candidate sequences are periodically decoded and evaluated using the original predictors. The final output is the highest-scoring feasible candidate encountered during optimization. Because $x_0$ is included in the candidate set, the procedure always returns a feasible sequence.

#### 7.6 Sampling and gradients in logit space

Each column $\Theta_{:,l}$ contains trainable preferences for the $A$ possible symbols at position $l$. For each of $K$ independent samples, we draw $G^k_{a,l} \sim \mathrm{Gumbel}(0,1)$ and compute

$$P^k_{a,l} = \frac{\exp((\Theta_{a,l} + G^k_{a,l})/\tau)}{\sum_{b=1}^{A} \exp((\Theta_{b,l} + G^k_{b,l})/\tau)}, \qquad x^k_{\mathrm{hard},:,l} = \mathrm{onehot}\Big(\arg\max_a P^k_{a,l}\Big)\,. \tag{4}$$

Thus, each Gumbel draw produces both a discrete sequence $x^k_{\text{hard}}$ and a soft counterpart $P^k$. The temperature $\tau$ controls the sharpness of the soft counterpart. We pass $x^k_{\text{STE}} = x^k_{\text{hard}} + P^k - \text{sg}(P^k)$ to each predictor, so its forward-pass score is evaluated on the discrete sequence, while its backward-pass gradient flows through $P^k$.

For either the target predictor or an auxiliary predictor, denoted generically by $f_q$, the score used in optimization is

$$\bar{f}_q(\Theta) = \frac{1}{K}\sum_{k=1}^{K} f_q(x^k_{\text{STE}}). \tag{5}$$

The chain rule and the straight-through estimator give the surrogate gradient for the preference of symbol $b$ at position $l$:

$$\begin{aligned}\frac{\partial \bar{f}_q}{\partial \Theta_{b,l}} &\overset{\text{STE}}{=} \frac{1}{K}\sum_{k=1}^{K}\sum_{a=1}^{A} \left.\frac{\partial f_q}{\partial x_{a,l}}\right|_{x^k_{\text{hard}}} \frac{\partial P^k_{a,l}}{\partial \Theta_{b,l}} \\ &= \frac{1}{K\tau}\sum_{k=1}^{K} P^k_{b,l}\left[\left.\frac{\partial f_q}{\partial x_{b,l}}\right|_{x^k_{\text{hard}}} - \sum_{a=1}^{A} P^k_{a,l}\left.\frac{\partial f_q}{\partial x_{a,l}}\right|_{x^k_{\text{hard}}}\right].\end{aligned} \tag{6}$$

Intuitively, the gradient compares the score's sensitivity to symbol $b$ with its probability-weighted sensitivity to the other symbols at that position. Applying this calculation to $f_T$ and each $f_{c_i}$ yields the target and constraint gradients used by GRACE. Because the scores are averaged before optimization, the constraints act on these sample-averaged predictions.

##### 7.7 First-order drift under damped projection

Let $d_\delta = P_\delta g_T$, where $P_\delta$ is defined in Eq.6 (main text), and write $A = J_c J_c^\top$. Substituting the definition of $P_\delta$ gives

$$\begin{aligned}J_c d_\delta &= \left[I - A(A+\delta I)^{-1}\right] J_c g_T \\ &= \delta (A+\delta I)^{-1} J_c g_T.\end{aligned} \tag{7}$$

Thus the directional derivatives generally do not vanish for $\delta > 0$. If $J_c$ has full row rank, $A$ is positive definite. Writing $\sigma_{\min}$ for the smallest singular value of $J_c$, we obtain

$$\|J_c d_\delta\|_2 \le \frac{\delta}{\sigma^2_{\min} + \delta}\|J_c g_T\|_2 < \|J_c g_T\|_2 \qquad \text{if } J_c g_T \neq 0. \tag{8}$$

The damped direction therefore reduces the *combined magnitude* of first-order constraint drift relative to the unprojected target gradient, but generally does not eliminate it. For a step of size $\eta$, the vector of constraint-property changes is

$$\bar{\boldsymbol{f}}_c(\Theta + \eta d_\delta) - \bar{\boldsymbol{f}}_c(\Theta) = \eta\,\delta(A+\delta I)^{-1} J_c g_T + O(\eta^2). \tag{9}$$

As $\delta \to 0$, Eq. (7) tends to zero, recovering the first-order preservation of exact null-space projection.

#### 8 Statistical Analysis

**Evaluation metric.** For each run, we report the improvement of the best feasible design over its starting sequence,

$$\Delta f_T = f_T(x^\star) - f_T(x_0), \tag{10}$$

in the task's native oracle units. If no feasible improvement is found, the starting sequence is returned and $\Delta f_T = 0$.

**Unit of inference.** The starting sequence. Where a task was run at more than one seed per start, the seeds are collapsed to their per-start median before any test, so $n$ counts starting sequences and not runs.

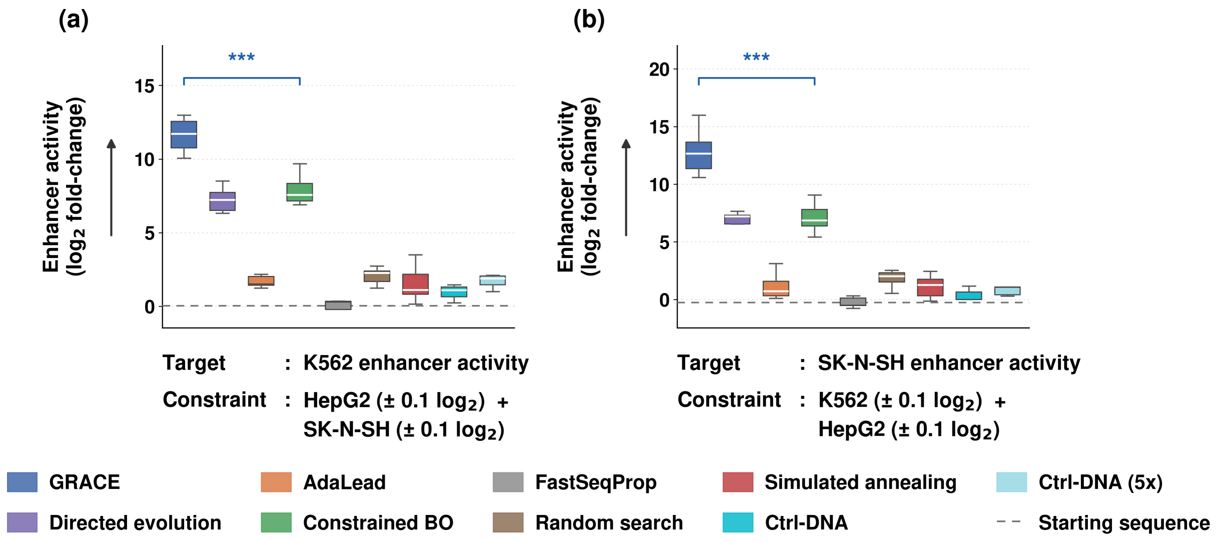

Figure 1: **Constrained enhancer optimization.** Improvements in predicted enhancer activity for (a) K562 and (b) SK-N-SH, while preserving activity in the other two cell types within $\pm 0.1$ $\log_2$ units. Boxplots show the distribution of best feasible improvements across runs. The dashed line indicates the starting activity ($\Delta f_T = 0$). $^{***}p < 0.001$.

**Statistical tests.** On each task GRACE is compared with the strongest baseline, the one with the highest median improvement. The comparison is paired by starting sequence and evaluated with a two-sided Wilcoxon signed-rank test. Markers are $^{*}p < 0.05$, $^{**}p < 0.01$, $^{***}p < 0.001$, and n.s. for $p \geq 0.05$. Tests are run separately per task and $p$-values are reported uncorrected.

**Sample sizes.** Ten tasks – the three enhancers, the three RNA tasks and the four CNN protein tasks – use 30 starting sequences at two seeds each, giving $n = 30$. Splicing uses 48 held-out splice sites, one run per site ($n = 48$). The two TEM-1 tasks share one reference sequence, so there the seed is the only available unit and $n = 8$. Heat shock uses 30 promoters and is the one task reported with restarts, described next.

**Heat-shock restarts.** A promoter is optimised several times from the same start under different random seeds and only the best feasible improvement is kept, so $n$ counts promoters rather than runs. Restarting is what one would do with a stochastic optimiser, and it matters here: about a third of single GRACE runs return the promoter unchanged, and the failures are seed-dependent rather than promoter-specific (promoter 6 gives 0.000 on seed 0 and 7.369 on seed 42). Coverage is uneven and the panel should be read with that in mind. GRACE has five seeds on all 30 promoters and AdaLead three to five on 20 of them, both at the matched 15,000-call budget; directed evolution is deterministic, so restarting leaves it unchanged. The remaining baselines come from an earlier step-matched set of 10 promoters at three seeds and are not collapsed over seeds. Paired tests use only the promoters that both methods ran.

### 9 Additional Results

Figure 1 presents additional results for cell-type-specific enhancer optimization. GRACE achieves the largest feasible improvements in both K562 and SK-N-SH activity, substantially outperforming the competing methods while preserving off-target activity within the prescribed tolerances.

Figure 2 provides two complementary redesign examples. GRACE improves predicted TEM-1 activity while preserving all eight annotated functional residues, and increases predicted 5′UTR ribosome loading while maintaining an open RNAfold structure (MFE = 0 kcal/mol).

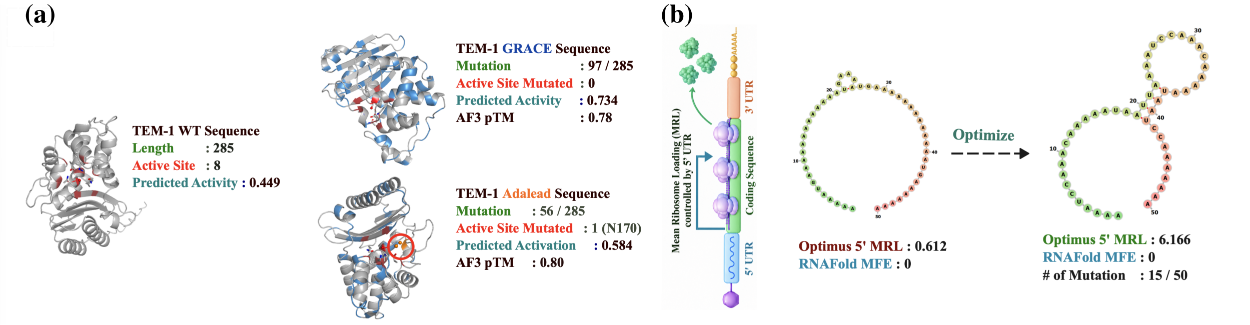

Figure 2: **Representative GRACE redesigns improve predicted function while preserving key properties.** **(a)** TEM-1 redesign. GRACE increases predicted activity from 0.449 to 0.734 while preserving all eight annotated catalytic and substrate-binding residues, despite 97 substitutions across the 285-residue protein. AdaLead reaches 0.584 and mutates the active-site residue N170. AlphaFold3 pTM scores are comparable for the GRACE and AdaLead designs (0.78 and 0.80). **(b)** A 50-nt 5′UTR redesign illustrating a goal relevant to mRNA vaccine expression: increase translation while keeping the 5′UTR open. The starting sequence has an unstructured predicted fold (RNAfold MFE = 0 kcal/mol) but low predicted mean ribosome loading (MRL = 0.612). With 15 substitutions, GRACE raises the Optimus 5-Prime predicted MRL to 6.166 while preserving an unstructured predicted fold (MFE = 0 kcal/mol). The resulting 5′UTR combines high predicted ribosome loading with the open structure sought for efficient translation.

### 10 Computational Resources and Runtime

Experiments were conducted on a Slurm cluster with hardware fixed across methods within each task. Protein CNN, RNA, and enhancer tasks used four CPU cores and 16 GB of memory; TEM-1 and heat-shock tasks used an A100 GPU, while splicing additionally used eight CPU cores. Runtime comparisons are therefore meaningful within tasks, rather than across different hardware configurations.

As shown in Fig. 3, GRACE requires only 7–25 s per run on CPU tasks, comparable to AdaLead (9–24 s) and FastSeqProp (14–28 s). In contrast, constrained Bayesian optimization requires 87–3619 s, approximately 10–200× longer than GRACE, primarily due to repeated surrogate fitting. Directed evolution and guided diffusion exhibit greater task-dependent runtime variation.

### 11 Reproducibility and Public Release

The benchmark and implementation will be released publicly. The release includes task definitions and constraint specifications, processed datasets with their train, validation, and test splits, the trained target and constraint oracles, the benchmark starting sequences, GRACE and all baseline implementations, evaluation and statistical-analysis scripts, and the configuration files needed to reproduce every reported number.

Three details are worth flagging for anyone reproducing the results, because each one silently changes the outcome if it is missed.

**Starting-sequence files.** Some task directories carry more than one candidate list of starting sequences from earlier curation rounds. Every result file records the constraint value $c_i^0$ of the sequence it actually started from, so the correct list can always be identified by matching those recorded values rather than by filename. Evaluation scripts in the release do this matching automatically.

**Genome build for the heat-shock library.** The MH4 fragment coordinates are on hg19. Extracting them against hg38 produces oracles that appear to train normally but reach Pearson 0.07 against measured activity. The released preprocessing pins the build explicitly.

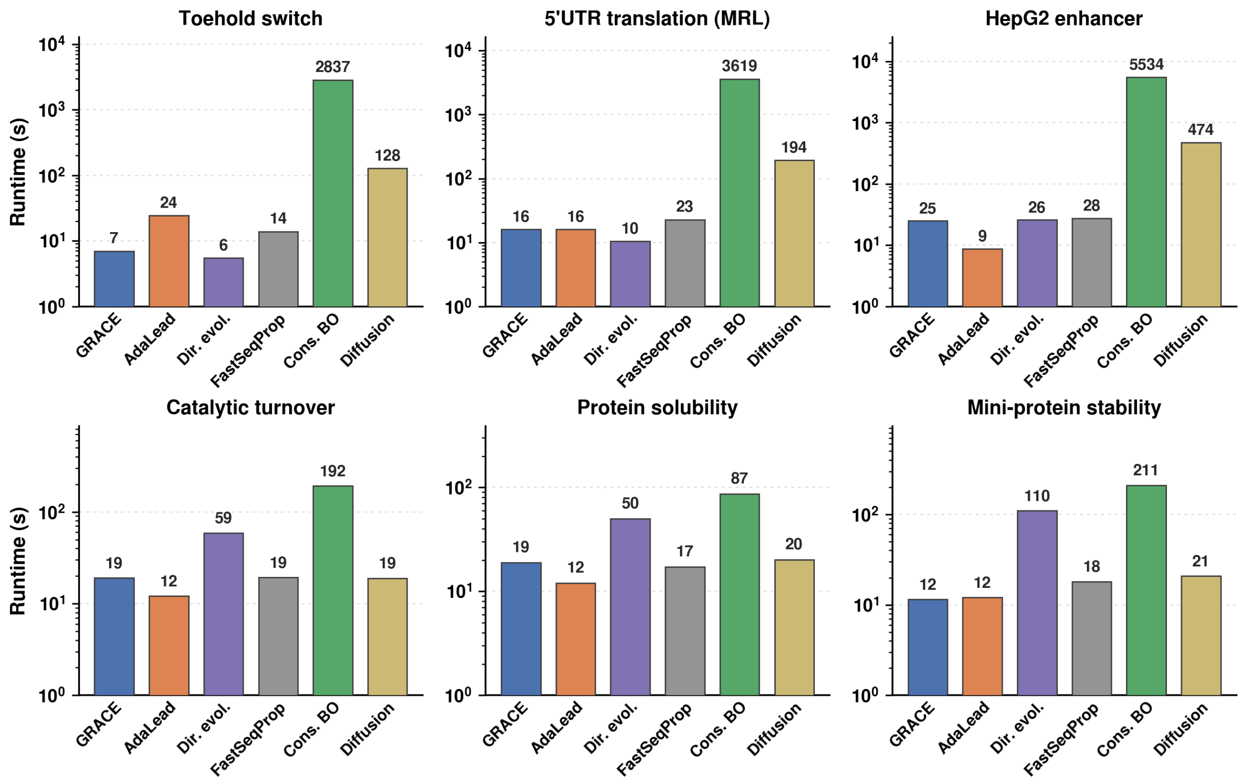

Figure 3: **Wall-clock runtime at matched predictor-call budgets.** Bars show one optimization run per method on task-specific hardware, with a logarithmic time axis.

**Evaluation-budget accounting.** Every result file records the number of oracle evaluations the run actually spent. Comparisons between methods should be made at matched counts of this quantity rather than at matched step or iteration counts, for the reasons given in Section 6.

**Repository.** The repository link will be provided in the camera-ready version.

[3] Maximilian Balandat, Brian Karrer, Daniel R. Jiang, Samuel Daulton, Benjamin Letham, Andrew Gordon Wilson, and Eytan Bakshy. BoTorch: A framework for efficient monte-carlo bayesian optimization. In *Advances in Neural Information Processing Systems*, volume 33, 2020.

[4] Lucía Barbadilla-Martínez, Noud Klaassen, Vinícius H. Franceschini-Santos, Jérémie Breda, Hatice Yücel, Miguel Hernández-Quiles, Tijs van Lieshout, Carlos G. Urzua Traslaviña, Minh Chau Luong Boi, Maryam Akbarzadeh, Celia Hermana-Garcia-Agullo, Sebastian Gregoricchio, Marcel de Haas, Roy Straver, Sarah Derks, Wilbert Zwart, Emile Voest, Lude Franke, Michiel Vermeulen, Jeroen de Ridder, and Bas van Steensel. Regulatory

grammar in human promoters uncovered by MPRA-based deep learning. *Nature*, 651:1107–1116, 2026. doi: 10.1038/s41586-025-10093-z.

[5] James Bergstra and Yoshua Bengio. Random search for hyper-parameter optimization. *Journal of Machine Learning Research*, 13:281–305, 2012.

[6] Xingyu Chen, Shihao Ma, Runsheng Lin, Jiecong Lin, and Bo Wang. Ctrl-dna: Controllable cell-type-specific regulatory dna design via constrained rl. *arXiv preprint arXiv:2505.20578*, 2025.

[7] Elad Firnberg, Jason W. Labonte, Jeffrey J. Gray, and Marc Ostermeier. A comprehensive, high-resolution map of a gene's fitness landscape. *Molecular Biology and Evolution*, 31(6):1581–1592, 2014. doi: 10.1093/molbev/msu081.

[8] Jacob Gardner, Matt Kusner, Zhixiang Xu, Kilian Weinberger, and John Cunningham. Bayesian optimization with inequality constraints. In *Proceedings of the 31st International Conference on Machine Learning*, volume 32 of *Proceedings of Machine Learning Research*, pages 937–945. PMLR, 2014.

[9] Sager J. Gosai, Rodrigo I. Castro, Natalia Fuentes, John C. Butts, et al. Machine-guided design of cell-type-targeting cis-regulatory elements. *Nature*, 634:1211–1220, 2024. doi: 10.1038/s41586-024-08070-z.

[10] Nate Gruver, Samuel Stanton, Nathan C. Frey, Tim G. J. Rudner, Isidro Hotzel, Julien Lafrance-Vanasse, Arvind Rajpal, Kyunghyun Cho, and Andrew Gordon Wilson. Protein design with guided discrete diffusion. In *Advances in Neural Information Processing Systems*, 2023.

[17] Roshan Rao, Nicholas Bhattacharya, Neil Thomas, Yan Duan, Xi Chen, John Canny, Pieter Abbeel, and Yun S. Song. Evaluating protein transfer learning with TAPE. In *Advances in Neural Information Processing Systems*, volume 32, 2019.

[18] Gabriel J. Rocklin, Tamuka M. Chidyausiku, Inna Goreshnik, Alex Ford, Scott Houliston, Alexander Lemak, Lauren Carter, Rashmi Ravichandran, Vikram K. Mulligan, Aaron Chevalier, Cheryl H. Arrowsmith, and David Baker. Global analysis of protein folding using massively parallel design, synthesis, and testing. *Science*, 357 (6347):168–175, 2017. doi: 10.1126/science.aan0693.

[19] Paul J. Sample, Ban Wang, David W. Reid, Vlad Presnyak, Iain McFadyen, David R. Morris, and Georg Seelig. Human 5' UTR design and variant effect prediction from a massively parallel translation assay. *Nature Biotechnology*, 37(7):803–809, 2019. doi: 10.1038/s41587-019-0164-5.

[20] Sam Sinai, Richard Wang, Alexander Whatley, Stewart Slocum, Elina Locane, and Eric D. Kelsic. AdaLead: A simple and robust adaptive greedy search algorithm for sequence design. *arXiv preprint arXiv:2010.02141*, 2020.

[21] Mike Thompson, Mariano Martín, Trinidad Sanmartín Olmo, Chandana Rajesh, Peter K. Koo, Benedetta Bolognesi, and Ben Lehner. Massive experimental quantification allows interpretable deep learning of protein aggregation. *Science Advances*, 11(18):eadt5111, 2025. doi: 10.1126/sciadv.adt5111.

[22] Tony Zeng and Yang I. Li. Predicting RNA splicing from DNA sequence using pangolin. *Genome Biology*, 23(1): 103, 2022. doi: 10.1186/s13059-022-02664-4.
